# An EpCAM/Trop2 mechanostat differentially regulates collective behaviour of human carcinoma cells

**DOI:** 10.1101/2022.10.03.510449

**Authors:** Azam Aslemarz, Marie Fagotto-Kaufmann, Artur Ruppel, Christine Fagotto-Kaufmann, Martial Balland, Paul Lasko, François Fagotto

## Abstract

EpCAM and its close relative Trop2 are well-known markers of carcinoma, but the potential role of these cell surface proteins in cancer metastasis remains unclear. They are known, however, to downregulate myosin-dependent contractility, a key parameter involved in cell adhesion and migration. We investigate here the morphogenetic impact of the high EpCAM and Trop2 levels typically found in epithelial breast cancer cells, using spheroids of MCF7 cells as an in vitro model. Intriguingly, EpCAM depletion stimulated spheroid cohesive spreading, while Trop2 depletion had the opposite effect. Combining cell biological and biophysical approaches, we demonstrate that while EpCAM and Trop2 both contribute to moderate cell contractility, their depletions differentially impact on the process of “wetting” a substrate, here both matrix and neighboring cells, by affecting the balance of cortical tension at cell and tissue interfaces. These distinct phenotypes can be explained by partial enrichment at specific interfaces. Differential distribution and antagonistic loss-of-function phenotypes are also observed in two other breast cancer cell lines with very different characteristics, suggesting that these are general properties of these two regulators. Our data are consistent with a simple model in which the EpCAM-Trop2 pair acts as a mechanostat that tunes local cortical tension, thus influencing a spectrum of adhesive and migratory behaviours.

## Introduction

Contractility of the actomyosin cellular cortex is a central determinant of cell and tissue adhesive and migratory properties. Understanding how contractility is controlled at the global and local scales is thus key to a mechanistic explanation of morphogenetic processes, including cancer metastasis. This is a complex question, because myosin-dependent contractility is implicated in multiple processes, that can be concurrent or even antagonistic. A multitude of regulators have been identified, and one important, long-standing question is how they can carry specific functions while all targeting the same molecular process.

In this context, EpCAM and Trop2 are particularly intriguing candidate regulators. EpCAM, also called TACSTD1, is a cell surface protein specifically expressed in epithelial tissues, and overexpressed in most human carcinomas (Rao et al., 2005; Went et al., 2004). It is an important cancer biomarker for diagnostic and therapeutic purposes (Gires and Bauerle, 2010). While the name EpCAM (Epithelial Cell Adhesion Molecule) was given based on its proposed function as a homotypic cell adhesion molecule (Balzar et al., 1999; Litvinov et al., 1994), EpCAM is currently viewed as a regulator of intracellular signalling, impacting on proliferation (Chaves-Perez et al., 2013; Maetzel et al., 2009) as well as on cell behaviour. The latter activity relies on its ability to downregulate myosin contractility (Barth et al., 2018; Fagotto, 2020a; Gaston et al., 2021; Maghzal et al., 2013; Salomon et al., 2017), providing an obvious putative link between EpCAM and cancer metastasis that remains to be established. While there is only one EpCAM in fish and amphibians, a second closely related gene, called Trop2 or TACSTD2 has appeared in reptilians, at the root of amniote evolution, as result of the duplication of the EpCAM gene. In humans, Trop2 is co-expressed with EpCAM in all epithelia, except in the intestine. Human EpCAM and Trop2 have high sequence similarity between themselves, and with EpCAM of lower vertebrates. Trop2 has been also linked to cancer, but, again, its actual impact on cell adhesion and migration, and on cancer invasion remains unclear (Fagotto and Aslemarz, 2020; Lenárt et al., 2020; Švec et al., 2022). EpCAM morphogenetic function is best understood in zebrafish and Xenopus embryos. In both systems, gain and loss-of-function (GOF and LOF) experiments showed that EpCAM acts positively on cell-cell adhesion and motility (Maghzal et al., 2010; Slanchev et al., 2009). In particular, EpCAM is required during gastrulation for dynamic tissue reorganization through cell-cell intercalation (Maghzal et al., 2010; Slanchev et al., 2009). At later stages of development, loss of EpCAM eventually caused severe defects in tissue integrity (Maghzal et al., 2013; Slanchev et al., 2009). We showed that all the EpCAM GOF and LOF embryonic phenotypes could be accounted for by its ability to downregulate myosin activity (Maghzal et al., 2013, 2010). At the molecular level, EpCAM directly binds and inhibits the novel class of PKC kinases, and by doing so it represses phosphorylation of myosin light chain (pMLC) by the PKD-Raf-Erk-MLCK pathway (Maghzal et al., 2013). Importantly, the loss of cadherin adhesion upon EpCAM LOF turned out to be a secondary effect of the acute upregulation of myosin contractility (Maghzal et al., 2013). Myosin repression through nPKC interaction appears conserved in human EpCAM and Trop2 (Maghzal et al., 2013). In human intestinal epithelial cells, where only EpCAM is expressed, its loss in a rare disorder called congenital tufting enteropathy (CTE) leads to disruption of the epithelium (Sivagnanam et al., 2008), due again to myosin overactivation and uncontrolled tension of the actomyosin cortex (Barth et al., 2018; Salomon et al., 2017). Note that a recent study on the migration of isolated intestinal Caco2 cells proposed an additional/alternative role of EpCAM in spatial regulation of RhoA and myosin activity via fast recycling endosomes (Gaston et al., 2021). The significance of this phenomenon in the context of the intact epithelium remains to be evaluated.

While the myosin repression by EpCAM appeared to favour adhesion and migration in the abovementioned models, it is likely to have different effects depending on the cell types and specific conditions, consistent with reported conflicting results on migration of human cell lines (reviewed in (Fagotto and Aslemarz, 2020). Also, while EpCAM and Trop2 are thought to be at least partly redundant (Fagotto and Aslemarz, 2020; Nakato et al., 2020; Szabo et al., 2022; Wu et al., 2020), the conservation of the two genes throughout amniotes argues for distinct functions, although direct experimental comparison at the cellular level has been missing so far. Thus, EpCAM and Trop2 constitute an intriguing case of two very similar myosin regulators, widely co-expressed in epithelia.

In an attempt to clarify their role in a cancer-relevant context, we chose breast epithelial MCF7 cells as a well-established model of EpCAM-positive carcinoma cells (Dai et al., 2017; Osta et al., 2004). MCF7 cells were particularly well suited because EpCAM and Trop2 expression is moderately high, representative of an average range found not only in breast, but also colon or lung cancer cell lines (Expression Atlas, https://www.ebi.ac.uk/gxa), and still in the order of magnitude of expression in normal epithelia. Furthermore, MCF7 cells display robust E-cadherin-mediated cell-cell adhesion as well as good migratory activity. EpCAM can be strongly downregulated in particular situations, such as induction of the ectodermal-mesenchymal transition (Pan et al., 2018; Sankpal et al., 2017; Vannier et al., 2013), justifying the physiological relevance of experimental acute manipulations. We thus analyzed the impact of changes in EpCAM and Trop2 levels on MCF7 spheroids spreading on a matrix as a model of dynamic rearrangement of a solid tissue, and we systematically characterized the effect on myosin contractility and adhesion, using a combination of cellular and biophysical approaches.

We confirmed that EpCAM and Trop2 both inhibit cortical myosin, but we made the unexpected discovery that they actually showed counter-balancing contributions to adhesion and collective migration. This could be explained by their partially differential localization to distinct cell interfaces, and the analysis of cellular and biophysical parameters highlighted how relatively modest differences in the balance of cellular tensions can deeply impact the morphogenetic outcome. Extension of the study to HCC1500 and MDA-MD-453, two other breast cancer lines with very different characteristics, indicated that differential localization and antagonistic effect on cell behaviour are general properties of the EpCAM/Trop2 pair.

## Results

### EpCAM depletion increases spreading of MCF7 spheroids and their cohesiveness, while Trop2 depletion has the opposite effect

In order to investigate the role of EpCAM and Trop2 in collective tissue behaviour, we used spheroids of breast cancer MCF7 cells as an in vitro model that mimics a solid tumor (Kramer et al., 2013). These spheroids were placed on a soft matrix of fibrillar collagen I as a physiologically relevant extracellular matrix (Insua-Rodríguez and Oskarsson, 2016). The collagen gel was about 20-50μm thick and sufficiently soft for spheroids to be capable to deform it (see convex ventral face of spheroids in Fig.1H’). Under these conditions, spheroids actively spread, which we typically imaged over 24hrs (Fig.1A, Movie 1). Spreading of a spheroid is an integrated process that involves adhesion to extracellular matrix and collective migration, together with the ability of the tissue mass to remain coherent while allowing cells to exchange neighbours by cell intercalation). EpCAM was depleted by transfection of siRNA one day before starting to form the spheroids, which were laid on collagen two days later. By that stage, EpCAM depletion was close to complete and remained so until the end of the assay (Fig.S1A,B). Spheroids formed from cells transfected with control siRNA spread efficiently, typically increasing their area by 3-to 4-fold (Fig.1A,E, Movie1). Note that the center of the spheroid tended to remain compact, while the periphery showed irregular contours, with cells or cell groups frequently sticking out of the cell mass (Fig.1A,H,H”). EpCAM knockdown (KD) resulted in a strong, highly reproducible increase in spreading (Fig.1B,E,H, Movie 2). After 24 hrs, spheroids had the shape of a flat pancake, only about 2-3 cells thick (Fig.1H’). Interestingly, these EpCAM KD spheroids expanded as a very cohesive sheet. While at high magnification numerous protrusions could be observed that emanated from the cells at the edge of the cell mass, similar to control spheroids (Fig.1H”), at a coarse grain view, this edge was overall strikingly smooth compared to control spheroids (Fig.1H). The distinctive morphologies of control and EpCAM KD, indicative of differences in tissue cohesiveness, were quantitatively expressed as “solidity”, a measurement that indicates how closely the surface can be described by a circle (Fig.1F). We validated the specificity of the KD phenotype by showing that it could be fully rescued by using MCF7 cells stably expressing a DOX-inducible EpCAM variant resistant to the siRNAs (Fig.S1G). In complementary GOF experiments, EpCAM overexpression decreased spheroid spreading, opposite to the KD phenotype (Fig.S1H). We also verified that the increased spreading of siEpCAM spheroids was not due to a higher proliferation rate by performing the spheroid assay in the presence of Mitomycin C (MMC), an anti-mitotic agent (Fig.S1F). We concluded that the increased spreading upon EpCAM KD was independent of cell proliferation.

**Figure 1.**
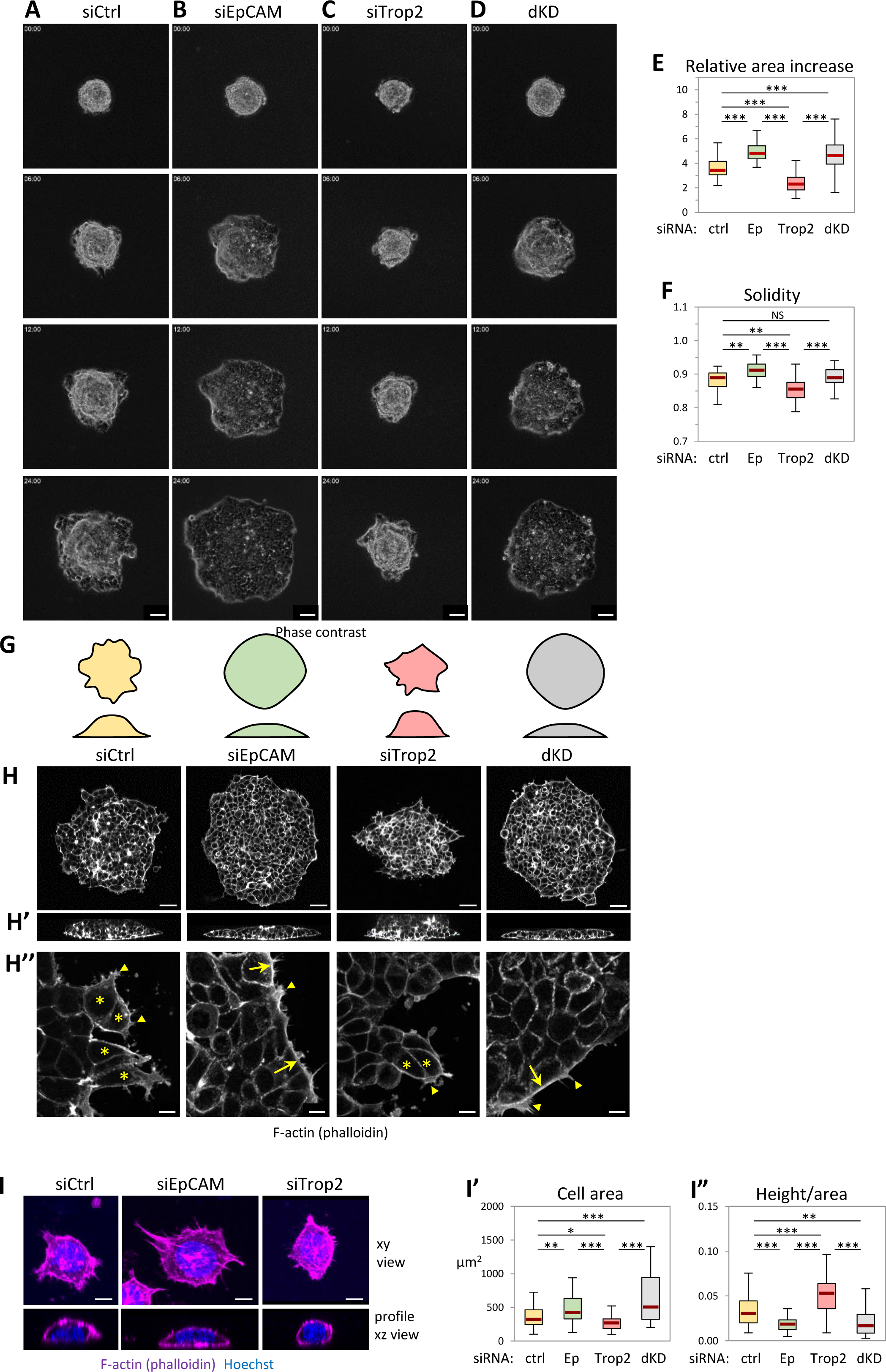
Cohesive collective migration of MCF7 spheroids is stimulated by EpCAM KD, but inhibited by Trop2 KD. Spheroids of transfected MCF7 cells with control, EpCAM, Trop2 and EpCAM/Trop2 siRNA were plated on a layer of fibrillar collagen gel, let adhere for 30 minutes, then phase contrast images were taken every 30min for 24hrs. **(A-D)** Images of whole spheroids at selected time points. Scale bar: 100µm. **(E)** Quantification of relative area increase after 24hrs, expressed as ratio of the final and initial areas. 45-63 spheroids from six to nine independent experiments, combining four experiments with siTrop2 sc-43043 and two with siTrop2 H502-0033, see supplemental FigS1E for separate results. In all figures, the box plots show the interquartile range (box limits), median (center line), and min and max values without outliers (whiskers). Statistical analysis: One-way non-parametric ANOVA (Kruskal-Wallis Test) followed by Dunn post hoc test. For all experiments presented in this study, P values are indicated as follows: * P < 0.05, ** P < 0.01, *** P < 0.001 and NS, not significant, only mentioned for pairs of interest. **(F)** Quantification of spheroids solidity, measured as the ratio [Area]/[Area of convex hull]. Quantification from the six experiments. **(G)** Schematic representation of typical spheroid morphologies in top and orthogonal views. **(H,H’)** Examples of phalloidin-labelled spheroids after 24hrs spreading in top and orthogonal view. Scale bars: 100μm. **(H”)** Details of spheroid edges. Maximal projection of 3 z planes, 1μm apart. Protrusions are observed in all conditions (arrowheads). However, control and Trop2 KD spheroids have numerous cells protruding out of the main cell mass (asterisks), which are rare in EpCAM KD and dKD. These are rather characterized by frequent actin cable-like structures along the spheroid edge (arrows). **(I)** Effect of EpCAM and Trop2 KD on single cell morphology. Typical examples of control, EpCAM and Trop2-depleted dissociated cells laid on collagen gel, stained with phalloidin and Hoechst. Top panels show maximal horizontal projection, the lower panel a slice from the orthogonal projection. Scale bars: 10μm. (**I’, I”)** Quantification of cell area, and calculation of height/area ratio. For each condition, 95, 82, 62 and 26 cells from three independent experiments. One-way ANOVA followed by Tukey-HSD post-hoc test.

Next, we depleted Trop2, using two distinct Trop2 siRNAs (sc-43043 and H502-003), which gave very similar results (Fig.S1C,D,E). For the rest of the study we used sc-43043. Using ectopic expression of EpCAM-GFP and Trop2-GFP to calibrate immunofluorescence (IF) signals, we estimated that the surface levels of EpCAM and Trop2 were quite similar (Fig.S2). Although Trop2 depletion was not as complete as for EpCAM (∼75% versus > 95%, Fig.S1A-D), phenotypes throughout this study were highly reproducible and consistent. Quite unexpectedly, Trop2 KD alone decreased spheroid spreading, opposite to the effect of EpCAM KD (Fig.1C,E,H, Movie 3). Transient expression of Trop2-GFP yielded a partial but significant rescue (Fig.S1I). The shape of Trop2 KD spheroids was strikingly irregular, yielding a low solidity value (Fig.1F). Thus, EpCAM and Trop2 appeared to play an antagonistic role in this context. We also examined double knockdowns for both EpCAM and Trop2 (dKD) and found that such spheroids did not spread more than EpCAM KD spheroids (Fig.1D,E,H, Fig.S1E, Movie 4).

The highly stereotypical morphologies adopted by spreading spheroids upon EpCAM KD and Trop2 KD was suggestive that these molecules acted on the “wetting” properties of the tissue (Fig.1G,H’). Indeed, the well-established biophysical analogy with the wetting properties of a liquid on a surface effectively accounts for the adhesive and spreading capacities of cells and tissues. They are based on the balance of “tensions” acting at the various cell interfaces (cell-matrix and cell-cell contacts, and free cell edges), which are largely dominated by cortical actomyosin contractility (Amack and Manning, 2012; Brodland, 2002; Douezan et al., 2011; Harris, 1976; Ryan et al., 2001; Steinberg, 1963; Winklbauer, 2015). Box 1 presents a summary of this biophysical principle, and introduces the concept of “adhesiveness” used in this study. Importantly, EpCAM and Trop2 KD not only influenced spreading by affecting adhesion to the matrix substrate, but apparently also by impacting tissue cohesion, which is related to cell-cell adhesion. Strikingly, we observed the exact same differences in morphology for single cells spreading on collagen gel (Fig.1I), as well as for small groups of cells (see below Fig.2G), indicating that a similar balance of tensions between free surface and substrate interface operated at the different scales of organization.

**Figure 2.**
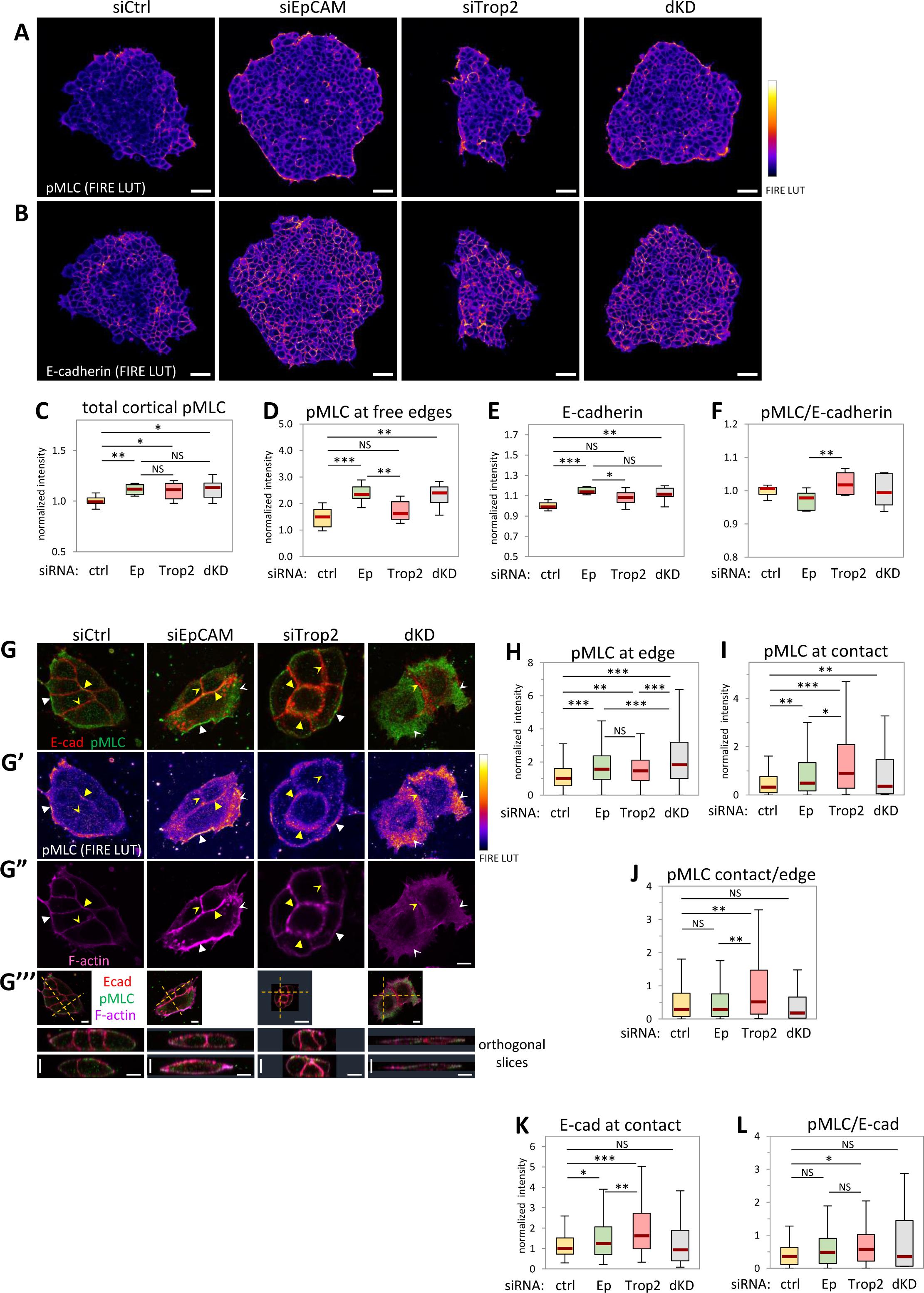
EpCAM and Trop2 depletions both enhance cortical myosin activation and E-cadherin recruitment, but differentially impact myosin at cell contacts and free edges. **(A,B) pMLC and E-cadherin levels in spheroids.** Representative confocal images of spheroids after 24hrs spreading, immunolabelled for p-MLC and E-cadherin. Levels are visualized using the “FIRE” pseudocolors (LUT) of ImageJ. The selected z planes correspond to the widest area for each spheroid. Scale bar: 50µm. **(C-F)** Quantification of normalized mean intensities for total p-MLC (C), pMLC along the free edge of the spheroid (D), and E-cadherin (E), and calculated pMLC to E-cadherin ratio (F). Both EpCAM and Trop2 KD cause an increase in both pMLC and E-cadherin. However, the pMLC to E-cadherin ratio is lowest in EpCAM KD but highest in Trop2 KD. Eight spheroids of each condition from three independent experiments. One-way non-parametric ANOVA (Kruskal-Wallis) followed by Dunn post hoc test. **(G-M) pMLC and E-cadherin levels in small cell groups. (G)** Representative confocal images of small groups of cells immunolabelled for E-cadherin (red) and pMLC (green), F-actin filaments labelled with phalloidin-Alexa647 (A”, magenta). Each image is the maximal projection of 5 planes, for a total thickness of 1μm. **(G’)** pMLC signal shown as FIRE LUT. Note that the image for siTrop2 is a slightly different plane as in A, to better visualize pMLC at contacts. White filled arrowheads: Signal along the free edge. Filled yellow arrowheads: Signal at cell contacts. Concave yellow arrowheads: Contacts with low to no pMLC signal. Concave white arrowheads: In very flat cells, the cortical signal is seen from the top (one edge of siEpCAM and the whole dKD pair). **(G’’’)** Comparison of size and shape through profile views (orthogonal slices along the dashed lines). Scale bars, horizontal and vertical: 10μm. **(H-M)** Quantification of peak intensities, determined through complete z stack series, for p-MLC along free cell edges (H) and cell-cell contacts (I), E-cadherin at contacts (K), F-actin at edges (M) and contacts (N). (J) Calculated ratio for pMLC between contacts and free edges. (L) Ratio pMLC/E-cadherin at contacts. pMLC and phalloidin intensities were normalized to the median value at edges of control cells. E-cadherin signal was normalized to control contacts. Data from seven independent experiments (∼80-180 cells/condition). One-way non-parametric ANOVA (Kruskal-Wallis Test) followed by Dunn’s post hoc test.

#### EpCAM and Trop2 depletions both lead to upregulation of myosin activity, but have distinct effects on local recruitment of myosin, cadherin and vinculin

Next, we characterized the effect of EpCAM and Trop2 depletions on the actomyosin cortex and adhesive structures. We conducted this analysis on whole spheroids, on small groups of cells, and on single cells in order to relate behaviour at the tissue scale to intrinsic cellular and biophysical properties.

We assessed the impact on myosin activity at the cell cortex by probing for phosphorylated myosin light chain (pMLC). Whole mount IF of spheroids showed that both EpCAM KD and Trop2 KD led to a mild but very consistent increase in global cortical pMLC (Fig.2A,C and Fig.S2D). We conclude that KD of either EpCAM or Trop2 indeed led to higher cortical myosin activity, consistent with their expected function. Beyond this general effect, we observed striking differences in pMLC pattern: In EpCAM KD, but not Trop2 KD, pMLC was enriched by ∼50% along the free edge of the spheroids (Fig.2A,D, Fig.S2D). Based on the principles presented in Box1, a higher increase in contractility at the edge relative to cell-cell contacts should correspond to increased cell-cell adhesion, and higher tissue cohesion. This is fully consistent with the measured smoothness of the circumference of siEpCAM spheroids (Fig.1F). On the contrary, in Trop2 KD, the increase in pMLC was biased toward cell contacts, suggesting lower cohesiveness, in agreement with the ragged outlines of these spheroids. Another striking effect of these depletions was an enhanced E-cadherin signal, quite significant in EpCAM KD, but much milder in Trop2 KD (Fig.2B,E). Cadherin recruitment upon increased tension is a hallmark of reinforcement of cadherin-dependent cell-cell adhesion (Box 1) (Engl et al., 2014; Gao et al., 2018), and cadherin levels are known to correlate with adhesiveness (David et al., 2014; Winklbauer, 2015). Importantly, the relative change in pMLC and E-cadherin, expressed as the total pMLC/E-cadherin ratio, was significantly lower in EpCAM KD than in Trop2 KD (Fig.2F), indicating that adhesion reinforcement was more efficient upon EpCAM KD..

We evaluated the same parameters on small groups of MCF7 cells (Fig.2G-L). The groups ranged from 2-6 cells, without noticeable differences within this size range. This range provided configurations where each cell in a group had at least one neighbour, was in contact with the matrix, and had a surface exposed to the medium (see orthogonal slice Fig.2G’’’). Since myosin and cadherins tend to concentrate at sites where most tension is exerted, we measured peak intensity for each interface, which gave a robust readout over the vast range of geometries observed for these cell groups. In controls, the pMLC signal was moderate at the free edges, and low, often undetectable at cell-cell contacts (Fig.2G’, concave yellow arrowheads), consistent with the well-established downregulation of tension along adhesive contacts (Winklbauer, 2015). The results were globally consistent with those of spheroids, despite some differences that were to be expected considering different cell configurations and readout. Firstly, cortical pMLC was increased both at edges and cell-cell contacts, confirming the shared role of the two molecules at downregulating contractility, but EpCAM KD had again a higher impact at edges relative to contacts, while the trend was opposite in Trop2 KD (Fig.2H,I). Thus, the signal ratio between contacts and edges was strongly increased in Trop2 KD, but not in EpCAM KD (Fig.2J). Both depletions also led to enhanced E-cadherin (Fig.2G,K). Here Trop2 KD yielded the most intense signal, but also again the highest pMLC/E-cadherin ratio (Fig.2L).

In summary, data from both spheroids and small cell groups largely coincided. They supported the common role of EpCAM and Trop2 in moderating cortical myosin, both at the free edge and at cell-cell contacts, consistent with the known function of Xenopus EpCAM (Maghzal et al., 2013). Furthermore, both depletions led to E-cadherin recruitment. Importantly, however, two potentially decisive differences emerged, a proportionally higher increase of pMLC at contacts compared to edges and a higher pMLC to E-cadherin ratio in Trop2 KD. Based on the principles enunciated above (Box 1), these observations were indicative of a fundamental difference in the balance of tensions, biased more toward cell contacts for Trop2 KD. As we will see below, measurements of tension and adhesiveness of cells confirmed this effect.

We next looked at the cell/tissue-matrix interface using vinculin, a classical marker of focal adhesions (FAs). Vinculin is recruited in a mechanosensitive manner, and its levels provide a readout for tension exerted on FAs. On a collagen gel, the ventral interface of spheroids was convex (Fig.1H’), reflecting the traction of the cells on the soft substrate. FAs could be accurately segmented for single cells (Fig.3A), but for cell groups, we analyzed cells seeded on a thinner layer of fibrillar collagen (∼ 5μm), where FAs could be clearly detected and segmented on a single horizontal plane (Fig.3B). On this substrate, cells appeared to sense the rigidity of the underlying glass, as witnessed by the strong ventral stress fibers (Fig.3B). The phenotypes for single cells on thick collagen and groups on thin collagen were in full agreement. For single cells, the total integrated FA signal per cell was increased in both EpCAM KD and Trop2 KD (Fig.3A’). However, when this signal was normalized relative to the ventral cell surface area (Fig.3A”), it appeared significantly higher for Trop2 KD, but not for EpCAM KD. FA vinculin intensity (per area) on large cell groups gave the exact same trend (Fig.3B’). These results clearly indicated that Trop2 KD led to a significantly more acute tension on the substrate that did EpCAM KD. Vinculin is also recruited in a tension-dependent manner at the cadherin-catenin complex (Leckband and de Rooij, 2014), and can therefore yield information on local tension at cell-cell adhesion sites. We thus analyzed cell contacts in the MCF7 monolayers on thin collagen (Fig.3B). Contacts between EpCAM-depleted cells had low vinculin, similar to controls. On the contrary, vinculin strongly accumulated at contacts of Trop2-depleted cells (Fig.3B”).

**Figure 3.**
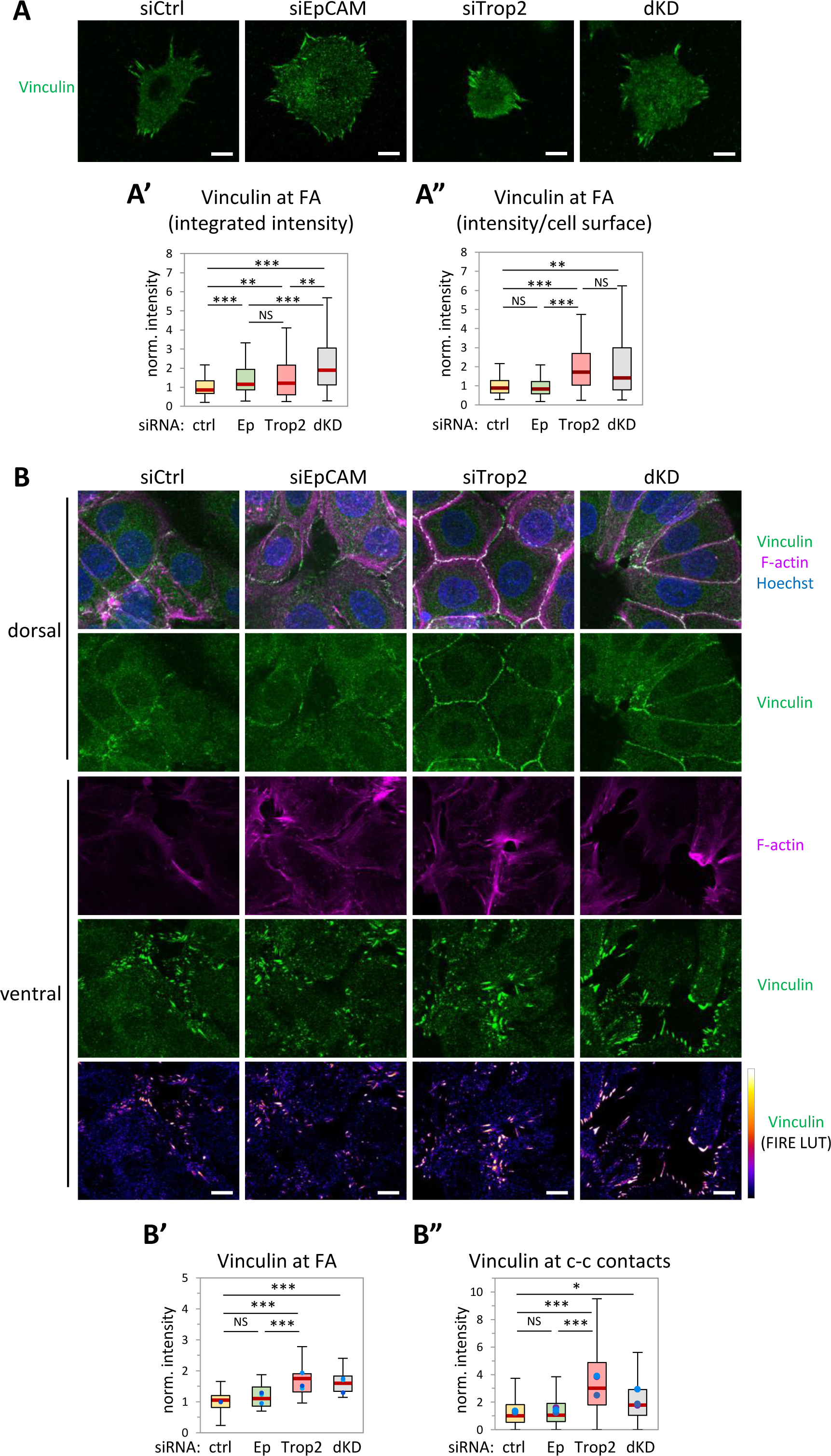
Different impact of EpCAM KD and Trop2 KD on recruitment of vinculin at focal adhesions and cell-cell contacts. **(A)** Confocal images of the ventral side (maximal projections) of single cells on collagen gel, immunolabelled for vinculin. **(A’)** Quantification of integrated vinculin focal adhesion intensity, normalized relative to the median value of control cells. **(A”)** Vinculin signal normalized to cell area. Data from six experiments (∼ 80 cells/condition). **(B)** Confocal images of large groups of cells laid on thin fibrillar collaen (∼5μm thick), immunolabelled for vinculin and stained with phalloidin. Maximal projections of deconvoluted images from z stacks, 1μm slices for the ventral side, ∼3μm slices for the dorsal side. **(B’)** Vinculin intensity at focal adhesions, normalized to average of controls, three experiments (18-20 fields/condition). **(B”)** Vinculin intensity along cell-cell contacts, three experiments (∼300 contacts/condition). Dots indicate medians of each experiment. A’,A”,B’B”: One-way non-parametric ANOVA (Kruskal-Wallis Test) followed by Dunn’s post hoc test. Scale bars: 10 μm.

For consistency, we also included dKD in all these experiments, although the effects were bound to be complex. dKD tended to give the highest intensities for total and edge pMLC (Fig.2C,D,H, only significant for the latter), and for total vinculin (Fig.3A), consistent with both EpCAM and Trop2 contributing to moderate cortical tension. Otherwise, dKD mostly gave intermediate values between those of the single KDs.

Taken together, the analyses of pMLC, E-cadherin and vinculin indicated that while depletion of either EpCAM or Trop2 boosted myosin activation at the cortex, there were key differences in their impact on cell-cell and cell-matrix interfaces, suggesting that tension at these adhesive interfaces increased more in Trop2 KD than EpCAM KD cells.

### Impact of EpCAM and Trop2 KD on cell-substrate and cell-cell forces

These observations prompted us to measure the actual forces exerted on different structures. We used traction force microscopy (TFM) on cell pairs adhering to H-shaped micropatterns (Fig.4A). The H patterns constrained the cell pair to adopt a stable configuration, providing a robust system to directly measure traction exerted on the matrix and, at the same time, to indirectly calculate the force at the cell-cell contact based on the balance of forces (Fig.4A’) (Tseng et al., 2012). In these settings, traction exerted on the matrix appears to be similar for single cells and for cell doublets (Ruppel et al., 2023). Another advantage was that the H pattern, by imposing a fixed geometry and a limited surface for the contacts to the matrix, reduced the wide differences in cell morphology of cells on non-constrained collagen substrate, naturally observed within the wild type MCF7 cell population, and further exacerbated by EpCAM and Trop2 depletions.

**Figure 4.**
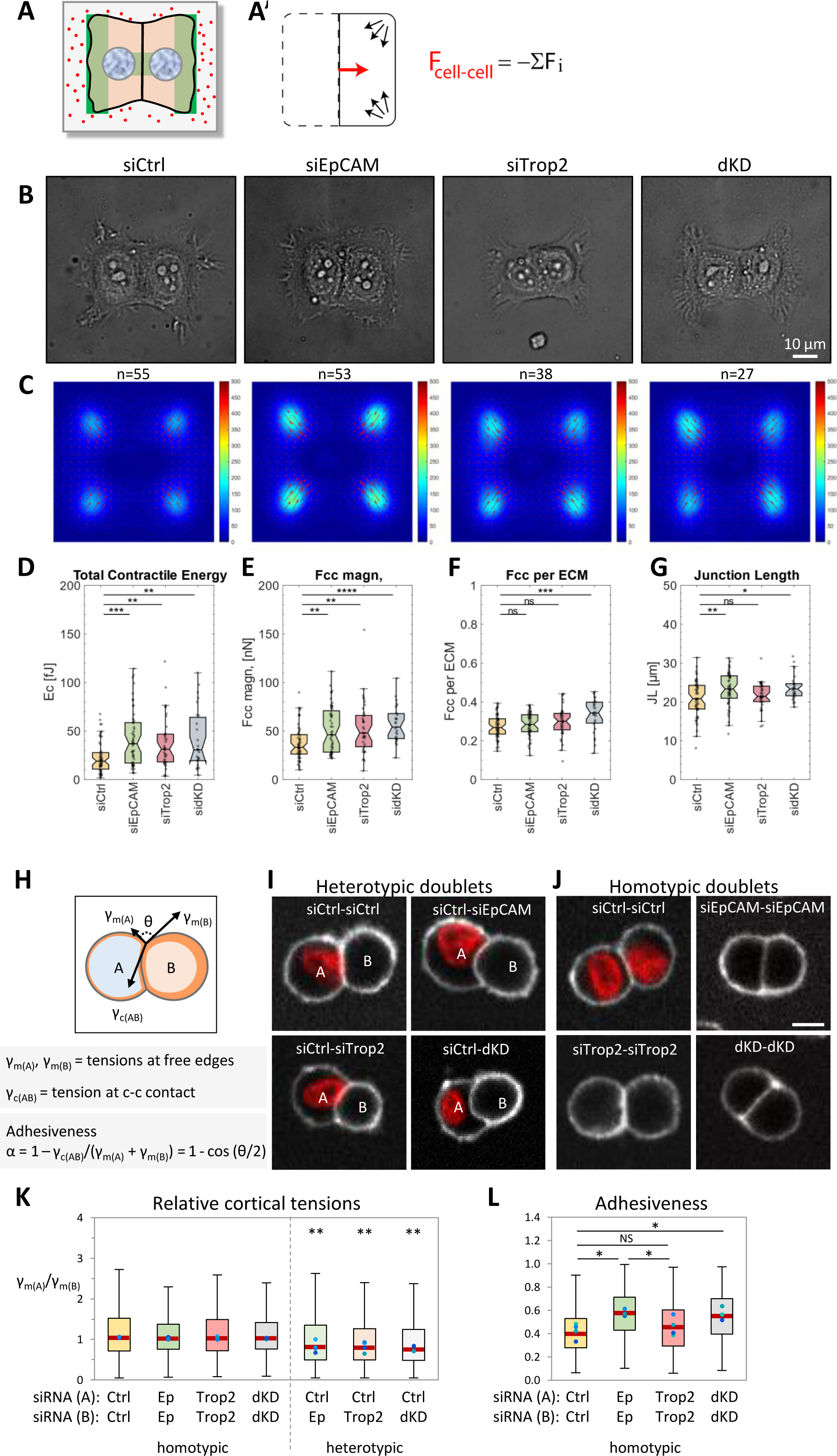
EpCAM KD and Trop2 KD differentially increase traction on substrate, cortical tension, and cell-cell adhesion. (A-D) Traction Force Microscopy (TFM) of cell doublets on H patterns. **(A)** Scheme of the experimental settings. Cells doublets were laid on H patterns coated with a thin layer of collagen (green) on a polyacrylamide gel with a stiffness of 5kPa, containing far red fluorescence nanobeads. Cells and underneath nanobeads were imaged, then cells were removed by trypsinization, and second images of the same positions were taken and used as reference, from which bead displacement was measured. **(A’)** Diagram of force vectors obtained by TFM on H-shape micropatterns. While traction forces (black arrows) are measured from the displacement of nanobeads, the cell-cell force (red arrow) is calculated indirectly based on it counterbalancing the sum of traction forces, as shown in the equation. **(B)** Representative images of micropattern-confined cell doublets for the four experimental conditions. **(C)** Corresponding average maps of traction forces. **(D)** Quantification of traction forces. **(E)** Quantification of cell-cell forces. **(F)** Cell-cell force to traction force ratio. **(G)** Quantification of junction lengths, measured from the phase contrast images. One-way ANOVA followed by Tukey-HSD post-hoc test. **(H-L) Determination of relative cortical tension and cell-cell adhesiveness of cell doublets laid on a non-adhering surface. (H)** Diagram of an asymmetric cell doublet, with the balance between cortical tensions at the free edges (γ_m(A)_ and γ_m(B)_) and contact tension (γ_c(AB)_) as introduced in Box 1. The principles of such system are as follows: The directions of the force vectors γ_m(A)_, γ_m(A)_ and γ_c(AB)_ are tangential to the membranes at the cell vertex, which allows to directly calculate the relative strengths of these tensions based on the geometry at vertices. The orange layer represents the actomyosin cortex, with its thickness symbolizing relative contractility. In the configuration represented in the diagram, the curved cell-cell interface reflects differences in cortical tension, as cell A tends to partly engulf B, due to its lower tension (γ_m(A)_ < γ_m(B)_). The strength of adhesion is related to the capacity of downregulating cortical tension along the cell-cell contact (Box 1). θ, the angle formed by the two vectors γ_m(A)_ and γ_m(B)_, directly relates to adhesiveness, a dimensionless value that ranges from 0 to 1. **(I,J)** Examples of homotypic and heterotypic doublets imaged by live confocal microscopy. The doublets were formed by mixing dissociated cells and let them re-associate on an adhesion-free support. Membranes were labeled with CellMask Alexa Fluor 647. siCtrl (A) cells were marked by Hoechst staining (showed in red) prior dissociation. Scale bar: 10µm. **(K)** Quantification of relative cortical tensions expressed as the ratio γ_m(A)_/γ_m(B)_. As expected, the ratio is close to 1 for homotypic doublets under all conditions. The ratio for heterotypic doublets is significantly lower, demonstrating that single and double depletions all cause an increase in cortical tension. **(L)** Quantification of adhesiveness for homotypic doublets, calculated from γ_c_ and γ_m_. Values show that EpCAM KD or dKD cells have increased adhesiveness compared to control cells. Trop2 KD only leads to a weak, non-significant increase. Four independent experiments, total 500-1250 doublets. Experiment averages indicated by dots. One-way non-parametric ANOVA (Kruskal-Wallis Test) followed by Dunn post-hoc test on experimental averages.

Using this approach, we found that the total tensile energy was increased in both EpCAM KD and Trop2 KD, reflecting a strong traction exerted on the FA (Fig.4C,D). These results directly demonstrated the increased contractility implied by the pMLC IF data. Yet, the length of cell-cell contacts, which is a readout of adhesiveness (see Box1 and below), was significantly broader for EpCAM KD pairs, while only marginally increased in Trop2 KD (Fig.4B,G). These results, fully consistent with the IF data, supported the notion that both EpCAM and Trop2 depletions upregulate myosin-dependent cortical tension, but adhesion reinforcement in Trop2 KD cells was less efficient. As expected, dKD also increased contractile energy, yielding the highest cell-cell force and highest cell-cell to total force ratio, consistent with EpCAM and Trop2 both contributing to moderating myosin contractility in these cells (Fig.4E,F).

The substrate used for TFM was an acrylamide gel coated with collagen. We verified the effect of EpCAM KD on spreading of spheroids on this acrylamide-based substrate. We found essentially the same phenotypes as on fibrillar collagen gel: EpCAM depletion strongly increased spreading, while maintaining high cohesiveness as indicated by high solidity values (Fig.S3A-D).

### Impact of EpCAM and Trop2 KD on cortical tension and cell-cell adhesiveness

An unequivocal method to evaluate differences in cortical tension and in cell-cell adhesiveness is to perform force inference based on the contact vertex geometry of free cell doublets on a non-adherent surface (Canty et al., 2017; David et al., 2014; Fagotto, 2020b; Parent et al., 2017; Winklbauer, 2015). Such doublets adopt a typical configuration, where the cell contact expands until contact tension γ_c_ is precisely balanced by the cortical tension at the interface with the medium γ_m_. As a consequence, a larger contact angle θ directly relates to higher adhesiveness (Box. 1). Moreover, if cell A and cell B have different cortical tensions γ_m(A)_ and γ_m(B)_, the heterotypic doublet is asymmetric, and the softer cell tends to engulf the stiffer cell (Fig.4H). Thus, the geometry of the contact vertex directly reflects the balance between the three tensions, γ_m(A)_, γ_m(B)_ and γ_c(AB)_ (Canty et al., 2017; David et al., 2014; Fagotto, 2020b; Kashkooli et al., 2021).

We adapted the protocol previously used for embryonic cells (Canty et al., 2017; Kashkooli et al., 2021; Rohani et al., 2014) to MCF7 monolayers for gentle dissociation into single cells. We then mixed at low density control and depleted populations, imaged both homotypic and heterotypic doublets, and determined the relative tensions (Fig.4I,J). These experiments led to the following conclusions: Firstly, heterotypic doublets made of a control (A) and a depleted cell (B) were clearly asymmetric (Fig.4I), resulting in γ_m(A)_/γ_m(B)_ significantly lower than 1 (Fig.4K). This constituted a firm demonstration that cortical contractility was indeed increased upon EpCAM and Trop2 KD. Second, contact angles θ of homotypic doublets were less acute for EpCAM KD than controls (Fig.4J), corresponding to a significant increase in adhesiveness (median 0.58 compared to 0.40 for controls (Fig.4L). In the case of Trop2 KD, however, only a small, non-significant increase was observed (Fig.4L). These two results were in perfect agreement with all our other data (spheroid morphometry, IF, TFM), demonstrating that while both KD led to increased contractility, this was accompanied by higher adhesiveness in the case EpCAM KD but not Trop2 KD.

### Spheroid spreading phenotypes can be simulated based on cell tensile properties

With this description of the differential impact of EpCAM and Trop2 depletions on the tensile and adhesive properties of the cells in hand, we asked whether these properties could account for the striking differences in the behaviour of the spheroids. With the principles enunciated in Box1, the experimental conditions can be described based on balance of the tensions exerted along the three cell interfaces γ_m_, γ_c_, and γ_x_. Cell-cell adhesiveness α is determined by the relative strengths of γ_c_ and of γ_m_ (Fig.4H, 5A-C). One key observation was that EpCAM KD cells had higher adhesiveness than control cells (Fig.4G,L), despite being more contractile (Fig.4D,K). We explained this effect by the fact that the increase was comparatively stronger for γ_m_ than γ_c_ (Fig.5A,B). The case of Trop2 KD was different: both γ_m_ and γ_c_ were also increased, but their ratio was roughly maintained, so adhesiveness did not significantly change (Fig.4G,L and Fig.5B). Spreading on the extracellular matrix can be similarly described by tensions γ_m_ and γ_x_, which are counterbalanced by the tension of the elastic substrate γ_s_ (for simplification, we consider here only the horizontal component) (Fig.5C). This balance of force can be expressed as γ_s_ = γ_x_ - γ_m_ x cos (Φ). As consequence of this relationship, if γ_s_ = γ_x_, the edge of the cell forms a right angle with the substrate. For γ_s_ - γ_x_ < 0, the cell spreads more on the substrate, forming an acute angle Φ, while for γ_s_ - γ_x_ > 0, Φ is obtuse, thus the cell tends to contract. These three situations corresponded to the conditions that were used in our experiments, respectively controls, EpCAM KD and Trop2 KD (Fig.5A,C). The correspondence was supported by the comparison between γ_s_ and the total contractile energies from TFM (Fig.4D), by the shape of single cells in xz projections (Fig.1I), and by the vinculin data (Fig.3), which were qualitatively consistent with relative γ_x_ values. It is essential here to note that the extent of spreading of a cell on the substrate (“wetting”) does not depend on absolute tension strengths, but on their relative values. Thus, while γ_m_, γ_x_ and γ_s_ were all increased in both EpCAM KD and Trop2 KD, the γ_x_ to γ_m_ ratio changed in opposite directions, causing EpCAM KD cells to be flatter, and Trop2 KD to be taller.

**Figure 5.**
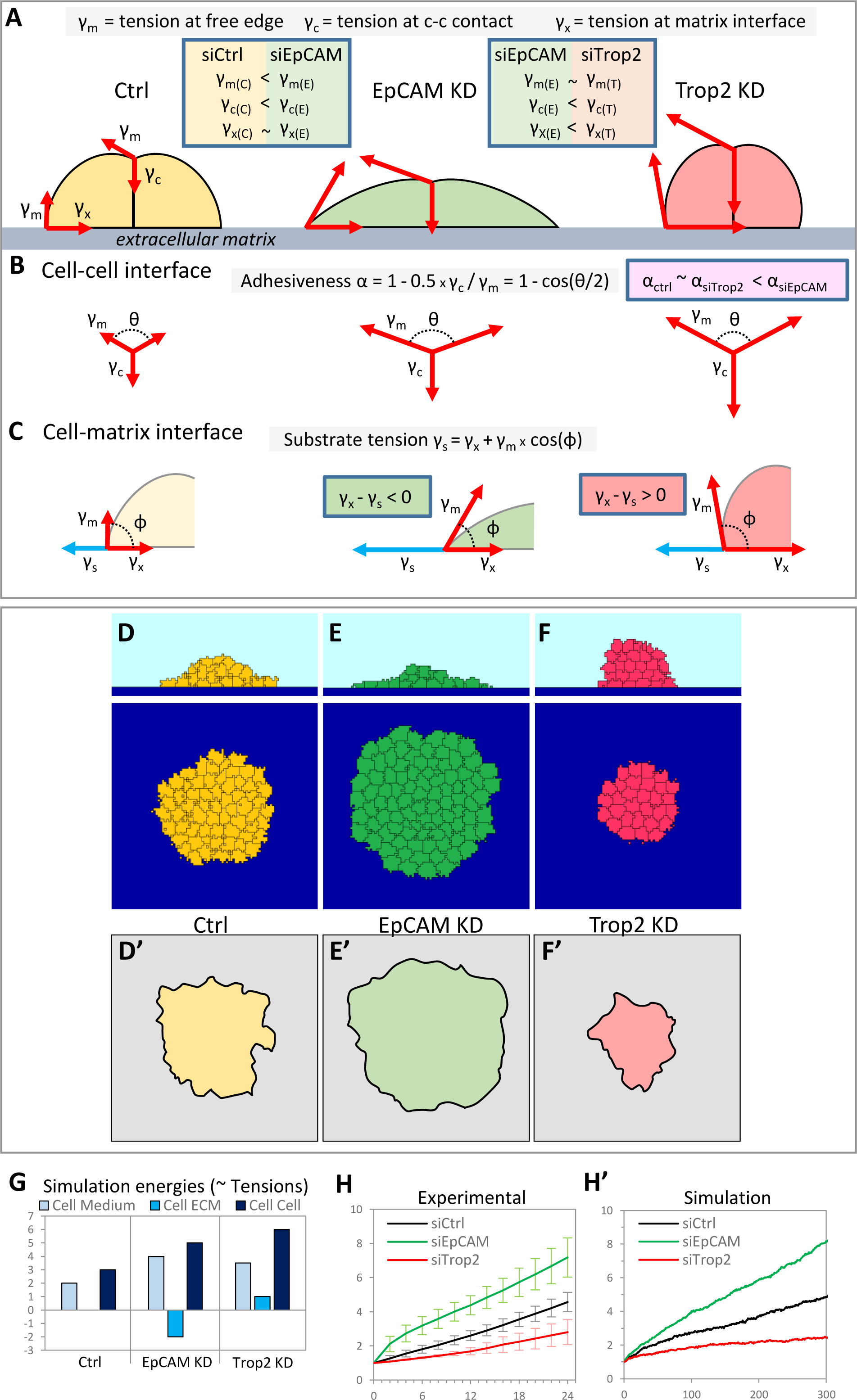
Simulation of MCF7 spheroid spreading in controls and under EpCAM/Trop2-depleted conditions based on biophysical parameters. **A.** Representation of schematic cell doublets on a substrate, with the relative tension vectors represented at the cell-cell vertex and at the edge of the matrix substrate interface. Their relative strength is represented by the length of the vector. The general changes in tensions between control and EpCAM KD, and between EpCAM KD and Trop2 KD are summarized in the two boxes with blue borders. **B**. Balance of forces at the cell-cell vertex. While all three tensions are strongly increased both in EpCAM KD and Trop2 KD, the resulting adhesiveness, which can be directly deduced from the angle θ, is increased in EpCAM KD but not in Trop2 KD. **C**. Balance of forces at the matrix interface. γ_s_ is the tension exerted on the substrate, corresponding to the traction measured by TFM. It counterbalances the cell-matrix tension γ_x_ and the horizontal component of the cell-medium tension γ_m_. Both tensions are increased in EpCAM and Trop2 KD, but their relative balance determines different outcomes: EpCAM KD cells adopts a spread configuration (acute angle Φ), while, on the contrary, Trop2 KD cells have a compact shape (obtuse angle Φ). For simplicity, only the horizontal component of the substrate tension is represented. **D-F.** Simulation of spheroids spreading using a 3D cellular Potts model (CompuCell3D software) for control, EpCAM KD and Trop2 KD conditions. See annex for detailed information. Images correspond to examples of yx and yz planes after 300 iterations. Medium is in light blue, matrix substrate in dark blue. **D’-D’.** Outlines of explants at 24hrs from Fig.1, drawn for comparison. **G.** Energy parameters used to simulate the three conditions. **H,H’.** Comparison of curves of relative area expansion from experimental data and from simulation. Error bars: SD.

With these cellular parameters at hands, we simulated spheroid spreading using a 3D version of a cellular Potts model (CompuCell3D (Cickovski et al., 2005; Izaguirre et al., 2004)). In this model, the “tensions” for a given cell type are input as relative “energies” (see appendix). While these energies are arbitrary values, they are related to γ_m_ for the cell-medium energy, γ_c_ for cell-cell energy, and γ_x_-γ_s_ for cell-matrix energy. We set the three energies (Fig.5G) to best fit both our biophysical data and the morphology of single cells and cell doublets (Fig.1I, S3E,F and 5A-C). The key features were (1) higher global energy for both EpCAM KD and Trop2 KD, which accounted for increased contractility compared to controls, (2) a lower cell-cell energy/cell-medium energy ratio (thus higher adhesiveness) for EpCAM KD, as well as (3) a cell-substrate energy set negatively for EpCAM KD (γ_x_-γ_s_ < 0) and positively for Trop2 KD (γ_x_-γ_s_ > 0). Simulation of the behaviour of spheroids under these three conditions yielded clear differences in the degree of spreading, moderate for the control condition, high for EpCAM KD, and low for Trop2 KD (Fig.5D-F, H, Movies 5 and 6). Thus, inputting parameters inferred from data on single cells/doublets faithfully reproduced the experimental spheroid spreading phenotypes (Fig.5D’-F’, H’). We concluded from these stimulations that the experimentally observed differences in cortical contractility at different interfaces were sufficient to account for the diametrically opposite spreading behaviours produced by EpCAM and Trop2 depletions.

### Partial differential distribution of EpCAM and Trop2 at cell interfaces may account for the distinct phenotypes

The opposite phenotypes of EpCAM and Trop2 KD could be explained by differences in distribution of the two regulators, which may then preferentially act on one or the other of the cell interfaces. We set up to determine these distributions by quantitative IF on small groups of cells. Consistent with their global impact on contractility and adhesiveness, EpCAM and Trop2 were both strongly expressed all along the plasma membrane, including cell-cell contacts. Standard IF with post-fixation permeabilization detected few EpCAM and/or Trop2-positive intracellular spots, that represented a negligible pool relative to the surface signal (Fig.S4E). Because permeabilization led to a significant loss of Trop2 signal at the plasma membrane, we present data from cell surface labelling without permeabilization, which gave robust signals for both proteins (Fig.6A). We quantified their distribution along the ventral side in cell-matrix contacts, the cell-cell contacts, as well as the dorsal side/lateral free edge, excluding protrusions that were measured separately (diagrams Fig.6D and S4A). These locations corresponded to the sites where the three basic cortical tensions γ_x_, γ_c_ and γ_m_ are exerted. While the two molecules showed a very similar general distribution (Fig.6A,S4B), we found a conspicuous quantitative difference between the free interface (dorsal side and lateral edges) and the ventral interface, the latter displaying proportionally more Trop2 than EpCAM (Fig.6A”,E). The resulting difference in dorsal/ventral ratio was modest (∼20%) but reproducible (Fig.6E). A more detailed analysis including additional subregions confirmed this (Fig.S4A-C). A second difference was observed specifically for cell-cell contacts at the interior of cell groups (here named inner contacts), where Trop2 was also enriched relative to EpCAM (Fig.6H,I). Loss-of-function situations led to a more contrasted landscape: Although EpCAM levels and distribution were largely unaffected in Trop2 KD cells, Trop2 was significantly impacted by EpCAM KD; its levels decreased on the whole lateral and dorsal free edges, while they increased at the ventral interface (Fig.S4C), resulting in a strong reduction in both the free edge/ventral ratio, and the free edge/cell-cell contact ratio (Fig.6F,G). These changes were of importance for interpreting the EpCAM KD phenotypes, since they were bound to exacerbate the tension imbalance caused by EpCAM loss, further increasing edge tension γ_m_, while mitigating the impact on contact tensions γ_x_ and γ_c_. This situation was perfectly consistent with EpCAM KD boosting spreading on the matrix as well as tissue cohesion. The case was opposite for Trop2 depletion, which was predicted to affect more acutely tension along the ventral interface and cell-cell contacts, explaining the decreased spreading and decreased tissue cohesion.

**Figure 6.**
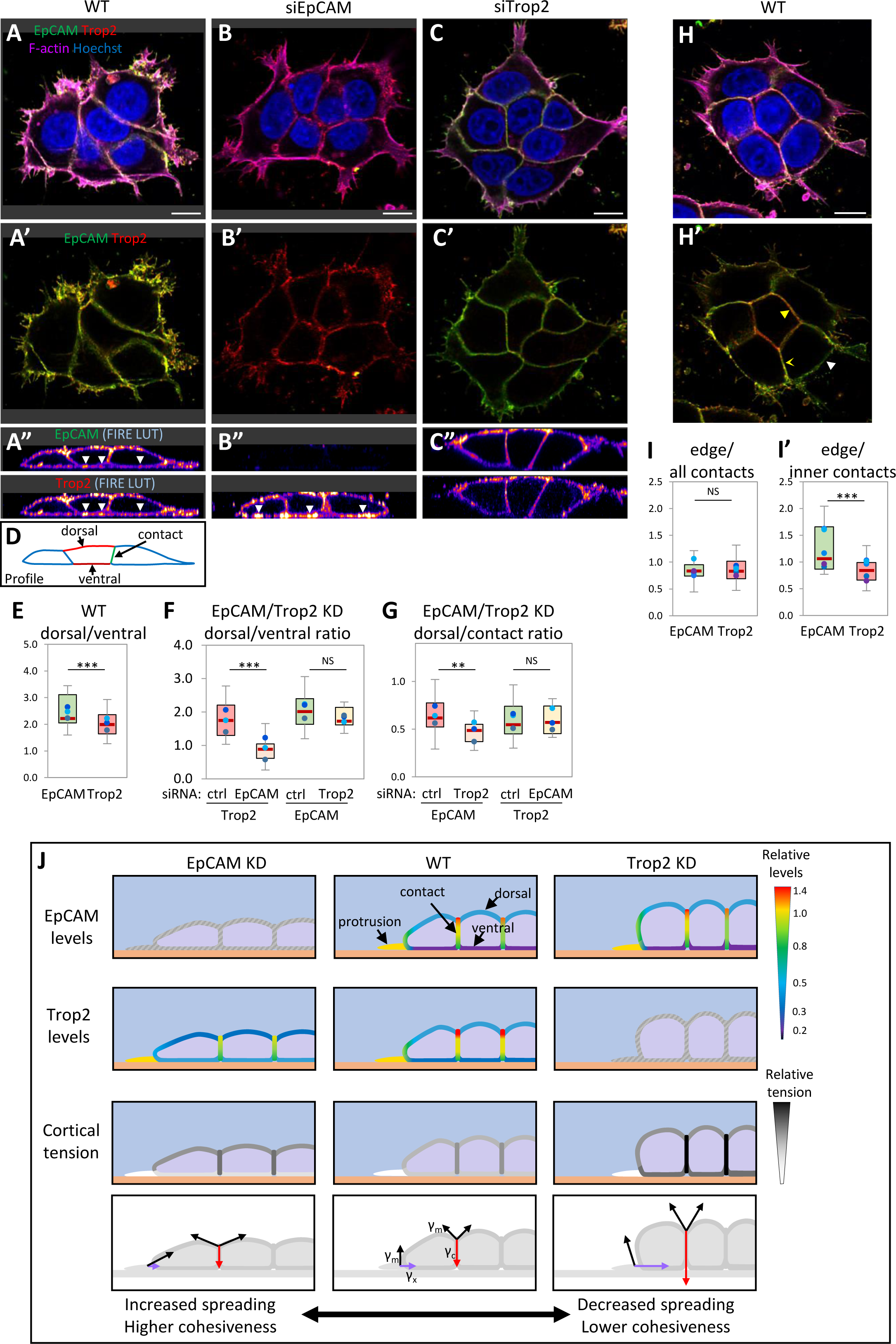
(A-I) EpCAM and Trop2 differential distribution along cell interfaces of MCF7 cells and impact of depletions. **(A-C)** Confocal microscopy image of small groups of wild type, EpCAM KD and Trop2 KD cells on collagen gel, labelled under non-permeabilized conditions for EpCAM (green), Trop2 (red) and F-actin (phalloidin-Alexa647, magenta). **(A,B,C)** Horizontal plane, merged image, maximal projection of 4 slices, 1μm total. **(A’,B’,C’)**. Merge of only green (EpCAM) and red (Trop2) channels. **(A’’,B’’,C’’)** Profile, orthogonal views of the same groups. EpCAM and Trop2 channels are displayed as “FIRE” LUT. Both EpCAM and Trop2 were found distributed all along the plasma membrane, but there was a slight but reproducible difference between the free edges (dorsal/lateral), richer in EpCAM, and the ventral side, richer in Trop2 (A”, white arrowheads). Note that Trop2 KD was incomplete (compare A” and C”). **(D-F)** Quantification. EpCAM and Trop2 levels where analyzed from whole z-stacks, as well from resliced stacks to obtain profile projections (see details in Fig.S4). **(D)** Diagram of the three main interfaces quantified based on profiles. **(E)** Pairwise comparison of dorsal and ventral interfaces for WT groups. **(F,G)** Dorsal/ventral and dorsal/contact ratios for controls and EpCAM or Trop2 KD. The ratios for EpCAM are not significantly affected by Trop2 KD, but the ratios for Trop2 are strongly decreased in EpCAM KD. See Fig.S4 for more detailed quantifications. Results from 13-14 cell groups, 3 experiments. Statistical analysis: Pairwise Student’s t-test. **(H)** Example of cell group with a central cell, surrounded by “inner” contacts (H,H’, yellow arrowheads). As obvious in the EpCAM/Trop2 merge of panel H’, these contacts have proportionally lower EpCAM/higher Trop2 compared to outer contacts (yellow concave arrowheads) and free edges (white arrowheads). **(I,I’)** Ratios edges/all contacts (I) and edges/inner contacts (I’), for EpCAM and Trop2. Quantification of 24 groups of 6-12 cells, from 5 experiments, of which 14 groups had inner contacts. Trop2 relative enrichment in inner contacts was also systematically observed in larger groups (example in Fig.S4D). Statistical analysis: Pairwise Student’s t-test. Scale bars: A-C, 5μm; H, 10μm. **(J) Summary diagram of EpCAM and Trop2 distribution in wild type and depleted MCF7, and impact on cortical tensions.** The schemes represent three cells at the edge of a group. The major regions are drawn, including dorsal, lateral and ventral surfaces, cell-cell contacts, and a leading edge lamellipodium. The collagen matrix is in orange. The two top rows of panels use a colour code to recapitulate the relative levels of EpCAM and of Trop2 under wild type and EpCAM/Trop2-depleted conditions. The third row provides a summary of cortical tension at the various interfaces using an arbitrary grey scale. The lower row represents the three corresponding cellular tensions γ_m_ (black), γ_c_ (red), and γ_x_ (purple).

In summary, EpCAM and Trop2 had a similarly broad distribution in wild type cells, consistent with both contributing to moderate global cortical contractility. Yet, partial asymmetries, further amplified in the case of EpCAM KD, could account for the changes in interfacial tensions estimated experimentally (Fig.4) and inputted in our simulations (Fig.5). The resulting tension imbalances could in turn explain the surprisingly opposite phenotypic outcomes in terms of spreading and collective migration.

### Differential LOF phenotypes and differential distribution of EpCAM and Trop2 in other types of breast cancer cells

We asked whether similar differences in EpCAM and Trop2 could also be detected in other cell lines. As mentioned in the introduction, we expected EpCAM and Trop2 LOF phenotypes to be highly context-dependent, yet the underlying logic may be conserved. We selected two additional luminal breast cancer cell lines, HCC1500 and MDA-MB-453, that express both EpCAM and Trop2 (The Human Protein Atlas), but differ in their morphologies and have behaviours departing in opposite directions from MCF7 cells: When grown on collagen, HCC1500 cells spread very poorly on matrix (Figs.7A,B,S5), but establish extensive cell-cell contacts and form coherent aggregates (Fig.7B,S5). On the contrary, MDA-MB-453 rapidly spread and migrate, and, while capable of cell-cell adhesion, form less extensive cell contacts (Fig.7D,S7).

**Figure 7.**
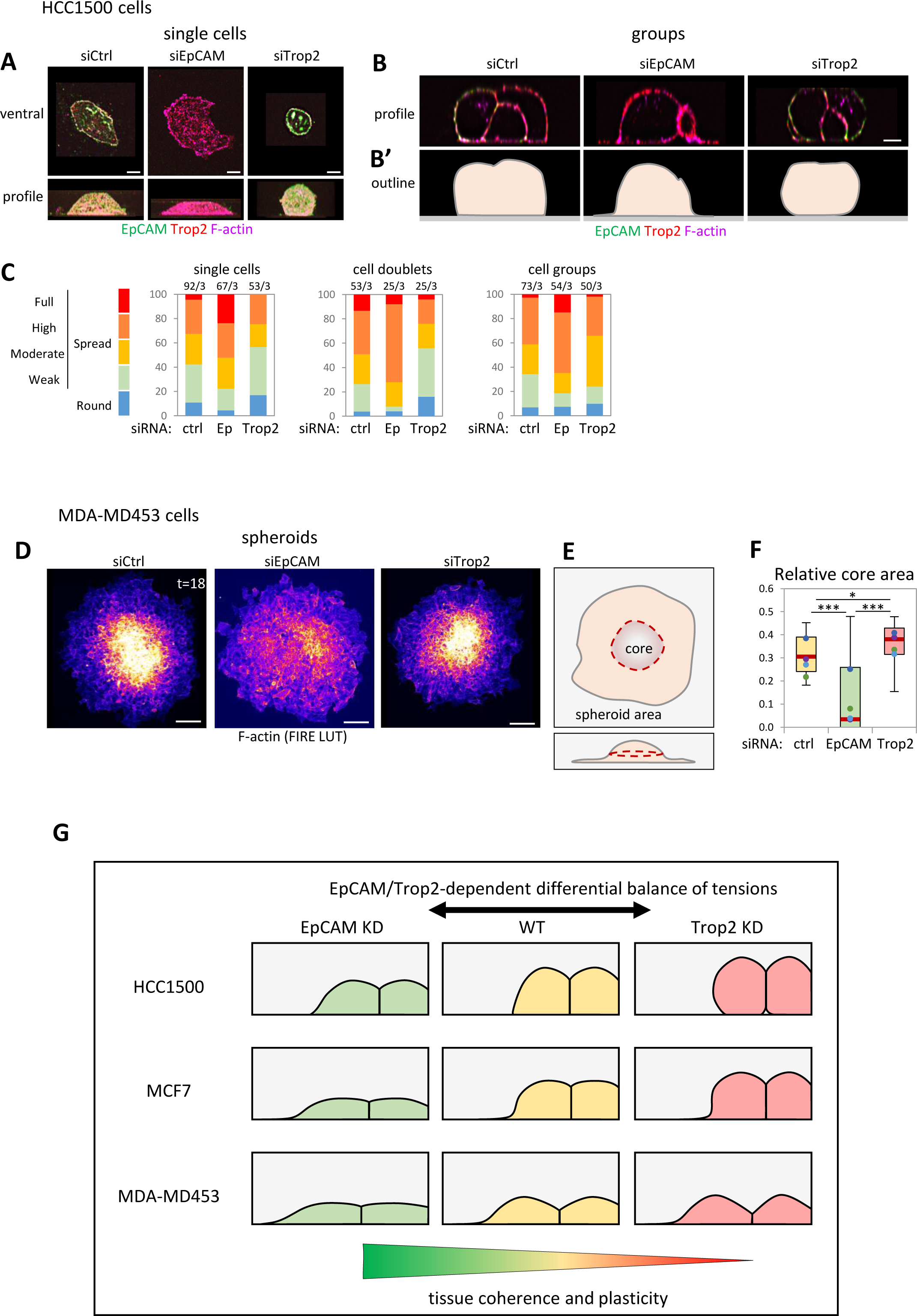
EpCAM KD and Trop2 KD phenotypes in other breast cancer cell lines. **(A-C) EpCAM KD and Trop2 KD in HCC1500 cells differentially impact on substrate wetting.** HCC1500 cells seeded on a collagen gel were cell surface immunolabelled for EpCAM and Trop2, stained with phalloidin, and imaged by confocal microscopy. The morphological analysis was made on profiles from orthogonal views, indicative of the degree of wetting, as detailed in supplemental Fig.S5. **(A,B)** Examples of single cells and groups in B. B’ highlight the outlines of the groups. Note that the extreme spreading observed for some single EpCAM KD cells shown in (A) was rarely found in cell groups. **(C)** Quantification. The bar graphs show compiled results for single cells, cell doublets and groups of cells, which all show similar trends. See Fig.S5D for statistical analysis. Scale bars, 5 μm. **(D-F) Distinct spheroid phenotypes of EpCAM KD and Trop2 KD in MDA-MD-453 cells. (D)** Control, EpCAM KD and Trop2 KD spheroids, 18hrs after adhering. Maximal projection of spread spheroids labelled for F-actin with Phalloidin, shown as FIRE LUT. See Fig.S7 for additional images, including z profiles. **(E)** Diagram indicating the typical central “dome” of a control spheroid. **(F)** Quantification of the ratio between dome and total area. The ratio was strongly decreased in EpCAM KD and increased in Trop2 KD. 30-33 spheroids from 4 experiments. One-way non-parametric ANOVA (Kruskal-Wallis Test) followed by Dunn post-hoc test. Bars, 50μm. **(G) Summary of EpCAM and Trop2 KD phenotypes in the three breast cancer cell lines tested in this study.** HCC1500, MCF7 and MDA-MD-453 are all luminal breast cancer cell lines that share in common the ability to form E-cadherin cell-cell contacts, but differ in their ability to spread on a collagen matrix (“wetting”). Cell-cell adhesion, thus tissue cohesion, appears to dominate over matrix adhesion for HCC1500, while for MDA-MD-453 the balance is tilted the other way. MCF7 represent an intermediate case. In all three cell lines, EpCAM KD tends to stimulate both spreading on the matrix AND cell-cell adhesion/tissue cohesion (flatter angles at contact vertices), while Trop2 KD tends on the contrary to increase tension at both types of interfaces.

EpCAM KD and Trop2 KD showed opposite phenotypes, in both cell types. Spreading of HCC1500 cells, single or in groups, was stimulated by EpCAM KD, but diminished upon Trop2 KD (Fig.7A,S5A,B). HCC1500 spheroids did not spread under any condition. However, we observed differences at cortical interfaces, monitored by phalloidin staining, with the intensity ratio between free edges and cell-cell contacts highest for EpCAM KD and lowest for Trop2 KD (Fig.S5E). Contrary to HCC1500, wild type MDA-MB-453 spheroids adhered and spread rapidly and more extensively than MCF7 spheroids (Fig.S7A). After 18hrs, a large portion of the cell mass had already migrated away from the original core, the remnant of which still emerged as a dome (Fig.7D,E,S7A,B). EpCAM or Trop2 KD did not significantly change global spheroid expansion (Fig.7D,S7A,C), but clearly impacted on spheroid morphology: Upon EpCAM KD, the spheroids spread into a flatter disc, with the central dome less prominent or absent (Fig.7D,F,S7B’). Furthermore, cells near the edge maintained tight contacts, suggesting that the spheroid expanded as a more coherent mass (Fig.S7B’). In Trop2 KD, the core of the spheroid remained more massive (Fig.7D,F,S7B’), surrounded by a loose monolayer of cells (Fig.7F,S7B’), suggesting that spheroid “spreading” was mostly due to individual cell migrating away, while the core failed to expand. Altogether, despite the large differences in basal characteristics of HCC1500 and MDA-MB-453, the effects observed for EpCAM KD and Trop2 KD were all consistent with a differential action at different interfaces, similar to MCF7 cells.

We determined the cell surface EpCAM and Trop2 distribution in wild type HCC1500 and MDA-MB-453 cells (Fig.S6,S7D,E). In HCC1500, the ratio between the dorsal and ventral sides was significantly higher for EpCAM than for Trop2, and the same trend was found comparing lateral edges to cell-cell contacts (Fig.S6). A similar dorsal to ventral difference was observed in MDA-MB-453 (Fig.S7D,E). We concluded that the partial asymmetric distribution of EpCAM and Trop2 between free edges and contact interfaces was shared at least by three very different breast cancer lines.

## Discussion

We uncovered in this study a subtle regulatory system that has a major impact on adhesive and migratory properties of cell aggregates. EpCAM and Trop2 are very closely related proteins, and we validated here that when expressed in the same cells, they both acted as negative regulators of myosin activity to moderate global cell contractility. This was evident from the quantification of cortical pMLC and firmly demonstrated by our TFM data. Nevertheless, depleting one or the other yielded to diametrically opposed phenotypes at the cell and tissue level, indicating that these closely related molecules function as a mechanostat in tissue morphogenesis. While we have discovered and characterized in detail this surprising phenomenon in MCF7 cells, we also presented confirming evidence in two other breast cancer cell lines, which suggests that this is a general property of this pair of molecules.

The most striking phenotype resulting from changing EpCAM or Trop2 levels is an opposite effect on spreading, consistently observed for single cells, cell groups and spheroids, and in all three cell lines. This effect is clearly linking EpCAM and Trop2 to a role in modulating substrate wetting (Douezan et al., 2011; Ryan et al., 2001), thus in acting on the balance of cortical tensions (Brodland, 2002). We should clarify at this point that actin polymerization-based protrusive activity also contributes to cell spreading and is certainly an essential component for cell migration. However, we consider this aspect as being at most peripheral to account for the EpCAM and Trop2-dependent phenotypes: In MCF7 cells, dynamic protrusions were observed in all situations and any degree of cell spreading, from flat EpCAM KD cells to round Trop2 KD cells. As for HCC1500 cells, most did not even form lamellipodia on collagen gel, and EpCAM depletion induced spreading of the whole cell “body”, generally in the absence of detectable “protrusions”, as evident from the z profiles. In parallel to matrix wetting, a second trait of the EpCAM and Trop2 KD phenotypes was the differential impact on cell-cell adhesion and tissue cohesion. Again, the same trend was found in the three different cell lines. As introduced in Box 1, cell adhesion is also a phenomenon of cell wetting, here cells maximizing their common interface. These observations amply support the view that cortical actomyosin contractility is the major target of EpCAM and Trop2 as far as cell behavior is concerned.

At first glance, the increased spreading and cohesion of EpCAM depleted MCF7 cells may seem at odds with the proadhesive role of EpCAM observed in lower vertebrate embryos and human intestinal cells (Barth et al., 2018; Maghzal et al., 2013, 2010; Salomon et al., 2017; Sivagnanam et al., 2008; Slanchev et al., 2009). We had predicted that the consequences on cell behaviour shall be context-dependent (Maghzal et al., 2013), contractility being on one side antagonistic to adhesion (Brodland, 2002), on the other side essential to stabilize and reinforce adhesion (Charras and Yap, 2018), a duality that considerably broadens the possible phenotypes (Box 1). However, the opposite EpCAM and Trop2 phenotypes in breast cancer cells came as a surprise. We propose that we are here dealing with two levels of regulation:

The first level concerns that capacity of adhesive structures to cope with increased cortical tension: In the case of Xenopus early embryos, cell-cell contacts appeared unable to resist to the strong myosin overactivation caused by EpCAM KD, resulting in lower adhesion, tissue stiffening and eventually disruption (Maghzal et al., 2013). In breast cancer cells, the increase in myosin activation and tension experienced upon depletion of either EpCAM, Trop2, or even both, was mild relative to wild type basal levels, and cells appeared to be able to cope with it though cadherin recruitment and adhesion reinforcement. This reaction was particularly efficient in EpCAM-depleted MCF7 spheroids, which managed the feat of strongly expanding while maintaining high tissue cohesion, and at the same time allowing extensive amount of intratissue cell movements (intercalation) required for their spreading.

The second level concerns the unexpected opposite phenotypes observed in cells expressing both EpCAM and Trop2. A simple model based on changes of the balance between the three tensions γ_m_, γ_c_ and γ_x_ provides an explanatory logic for all the observed phenotypes. An important feature of this model is that relatively small variations can have a large effect on the final behavioral output. Modulation of this balance can in turn be accounted for by partial differential distribution of the two negative regulators, combined with a dependency of Trop2 localization on EpCAM. Comparison of the three cell lines offers a preliminary glance at the spectrum of behaviours that can be achieved by the combined activity of the two partly concurrent, partly antagonistic regulators (Fig.7G). We propose that EpCAM and Trop2 modulate cell behaviour without necessarily overriding the intrinsic cell characteristics. Wild type HCC1500 cells, for instance, spread very poorly on collagen, and, while they did spread slightly better upon EpCAM KD, they were definitely not transformed into a mesenchymal-type cell. Conversely, in the case of the much less coherent MDA-MD-453 cells, EpCAM KD or Trop2 KD influenced spheroid compaction, but did not significantly affect their fast expansion. EpCAM and Trop2 LOF phenotypes were most spectacular in MCF7 cells, presumably because these cells are in a regime which is close to a turning point, where moderate changes in tension can significantly tilt cell and tissue behaviour.

While we find quite enticing that such spectacular phenotypes may be explained by small differences in distribution impacting on the local balance of cortical contractility, we should not exclude other alternate or complementary possibilities, including differences in biochemical activities of the two proteins. At the moment, myosin downregulation is the only firmly validated property related to morphogenetic properties, but future studies may provide additional mechanisms. Furthermore, the anti-myosin activity may not necessarily be uniform over the various cell interfaces, but different pools may exist. A first hint is the more punctate IF signal for Trop2 and its sensitivity to detergents, although no clear correlation could be inferred. Solving the details of this system, including the molecular basis of differential localization, will not be straightforward: All properties reported for EpCAM appear to be shared by Trop2, including oligomerization, association with tetraspanin-based domains, with claudins, potential interaction with the cytoskeleton, or matriptase-mediated cleavage (Balzar et al., 2001, 1998; Fagotto and Aslemarz, 2020; Guerra et al., 2022; Kuhn et al., 2007; Nubel et al., 2009; Pavsic et al., 2014; Pavšič, 2021; Szabo et al., 2022; Wu et al., 2020). Both form homodimers, but structural considerations indicate incompatibility for heterodimers (Pavšič, 2021). A related puzzling phenomenon is the sensitivity of Trop2 to EpCAM depletion, both in terms of level and subcellular localization, indicating a complex relationship, probably involving both cooperation and competition for recruitment to sub-compartments and/or for targeting to degradative routes.

How can one reconcile our results with the apparent redundancy indicated by the normal embryonic development of single knock-out mice, and with the partial compensation observed in tissues where both are expressed (Guerra et al., 2012; Lei et al., 2012; Szabo et al., 2022; Wang et al., 2011)? One piece of explanation may relate to the acute LOFs used in our study (over 2-5 days). It could be conceivable that, on the longer term, cells adapt to this loss by rerouting the remaining partner. We propose that, in the absence of compensatory mechanisms, EpCAM and Trop2 may be interchangeable for their generic action on contractility, but not for more refined balance of tension required in particular tissues. This is supported in the case of human CTE, where intestinal tuft formation caused by mutations in EpCAM was only partially rescued by ectopic Trop2 expression (Nakato et al., 2020).

While we are well aware of the caveats inherent to any LOF/GOF experiments, we consider that the depletion/overexpression approach used in this study is of clear physiological relevance, since the magnitude of the changes in EpCAM and Trop2 levels is compatible with what happens during cancer development. MCF7 cells represent in this regard a good model, based on the elevated expression levels compared to normal epithelia as well as on the abrupt drop in EpCAM expression observed after EMT (Sankpal et al., 2017). While this study has only addressed the function of EpCAM and Trop2 in three cell lines in an *in vitro* context, we hypothesize that co-expression of EpCAM and Trop2 may act as a general mechanostat contributing to setting the adhesive and migratory properties, which may be broadly relevant in the context of the biology of carcinoma. Significant levels of both EpCAM and Trop2 are found in a large number of cancer cell lines and multiple types of human cancers (Expression Atlas, https://www.ebi.ac.uk/gxa/ and Protein Atlas https://www.proteinatlas.org/). Our data may contribute to explain why one cannot assign a general unambiguous pro- or anti-invasive activity to these molecules. Indeed, their action shall not only depend on expression levels, but crucially on their relative levels, and will be further strongly influence by other, cell type-dependent properties, as exemplified in our study. One may reasonably predict that cancer cells can modulate this mechanostat, resulting in more plasticity during various phases of cancer development.

## Materials and methods

### Cell culture

MCF-7, HCC1500 and MDA-MD-453 cells, originally acquired from ATCC, were provided by the SIRIC Montpellier center, courtesy of Dr. Marion Lapierre. They were grown in complete culture medium, i.e. Dulbecco’s modified Eagle’s medium (DMEM) with 4.5 g/l glucose, supplemented with 10% fetal bovine serum (FBS), 1% antibiotic-antimycotic and 1% non-essential amino acids (all Gibco/Thermo Fischer Scientific), at 37°C and 5% CO_2_, for a maximum of 10 passages. 0.05% Trypsin-EDTA (Gibco/Thermo Fischer Scientific) was used for dissociation and passage.

### Antibodies, reagents, and solutions

The list of antibodies and dilutions used for IF are presented in Table 1. Secondary antibodies and phalloidin conjugated to Alexa fluorophores were from Molecular Probes/Thermo Fischer Scientific. EpCAM (sc-43032), Trop2 (sc-72392) and control non-targeting (sc-37007) siRNA were from Santa Cruz Biotechnology. A second Trop2 siRNA (H502-0033) was obtained from Sigma/Merck. Mitomycin C was from Sigma/Merck (M4287). Calphostin and Blebbistatin from EMD/Millipore. The EdU click reaction kit with Alexa Fluor 488 dye was from Invitrogen/Thermo Fisher Scientific (C10337). Phosphate Buffer Saline (PBS) without calcium and magnesium was from EuroBio (CS1PBS01-01). Tris Buffer Saline (TBS) was 100mM Tris-HCl pH 7.2, 150mM NaCl. Dissociation buffer was 88mM NaCl, 1mM KCl and 10mM NaHCO3, pH = 9.5 (Canty et al., 2017).

**Table 1.**
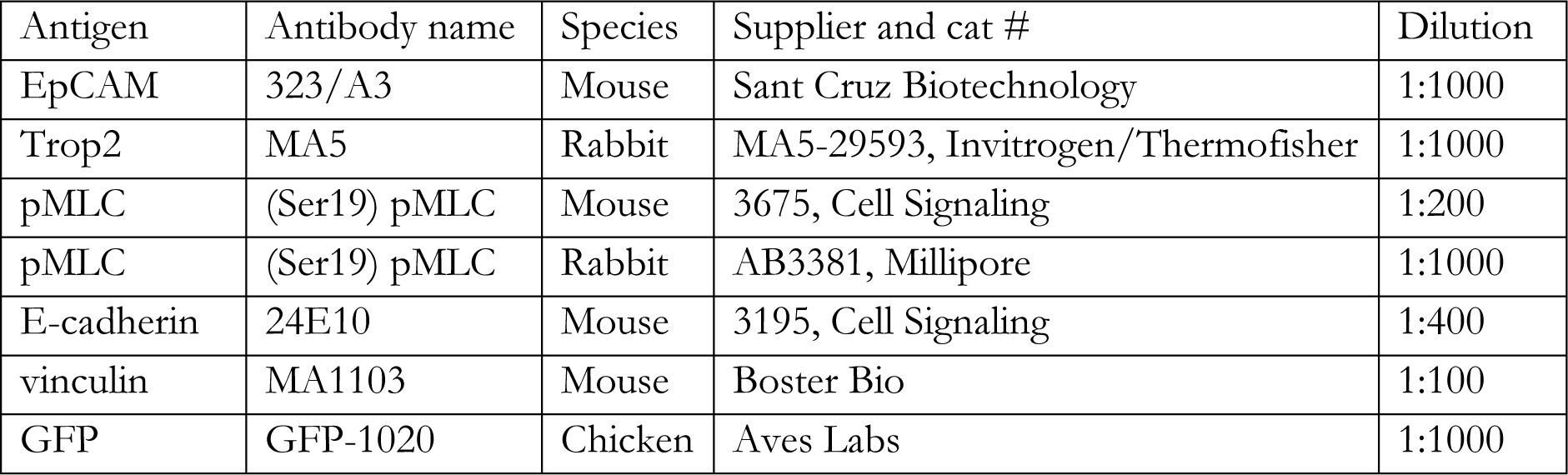
List of antibodies.

| Antigen | Antibody name | Species | Supplier and cat # | Dilution |
| --- | --- | --- | --- | --- |
| EpCAM | 323/A3 | Mouse | Sant Cruz Biotechnology | 1:1000 |
| Trop2 | MA5 | Rabbit | MA5-29593, Invitrogen/Thermofisher | 1:1000 |
| pMLC | (Ser19) pMLC | Mouse | 3675, Cell Signaling | 1:200 |
| pMLC | (Ser19) pMLC | Rabbit | AB3381, Millipore | 1:1000 |
| E-cadherin | 24E10 | Mouse | 3195, Cell Signaling | 1:400 |
| vinculin | MA1103 | Mouse | Boster Bio | 1:100 |
| GFP | GFP-1020 | Chicken | Aves Labs | 1:1000 |

### siRNA transfection

Cells were seeded in 24-well plates at 0.2 × 10^6^ density one day before siRNA transfection, and 1.6ml fresh medium was added just before transfection. 40 picomoles siRNA (2x 40 picomoles for dKD) and of 4μl Lipofectamine RNAiMAX (8μl for kD) (Invitrogen/Thermo Fischer Scientific) were added to each well. Efficient KD was obtained after 48 to 72 hours.

### Fibrillar collagen gels

Glass bottom dishes (Cellvis, D35-20-1.5-N) or 12mm round coverslips were plasma cleaned to create a hydrophilic surface, then coated at 4°C with a thin layer of a 2mg/ml solution of collagen I (from a ∼3mg/ml stock solution in 0.02N acetic acid) in PBS, titrated to alkalinity by addition of NaOH, and incubated at 37 °C for 30 min for gelling.

### Spheroid formation

The siRNA transfected cells were dissociated by trypsin and counted using trypan-blue solution and an automated cell counter (Invitrogen, Countess). For each spheroid, 400 cells were suspended in 200 μl complete culture medium containing 20% methylcellulose (Sigma, M0262-100G) and incubated in single wells of round-bottom 96-well plates (Greiner Bio-One Cellstar, 650185) for 48hrs.

### Spheroid migration assay

The formed spheroids were transferred to glass-bottom dishes coated with collagen gel. Adherence to the gel was observed within 30 min. Spheroids were imaged with an Olympus IX83 inverted widefield video-microscope controlled by Metamorph software using a 10x (0.3NA) or 20x (0.45NA) objective. Time lapse images were acquired every 30 min for 24hrs.

### Single cell migration assay

Cells were dissociated in dissociation buffer and counted as above, 0.5 × 10^5^ cells were plated on top of collage gel and left to adhere for about 3 hours before acquisition of time lapses. Imaging conditions were as for spheroids, with a 20x (0.45NA) objective, every 5 to 7 min for 15 hours. Migration was measured using the MetaMorph software tracking tool.

### Immunostaining

Spheroids (in glass-bottom dishes) were fixed for 30 min in 3.7% paraformaldehyde (PFA, #15714, Electron Microscopy Sciences) in PHEM buffer (60mM PIPES, 25mM HEPES, 10mM EGTA, and 4mM MgSO4, pH 7.2), followed by 30 min permeabilization in 1% Triton X100 in PBS. Fixed samples were washed twice with TBS, incubated for 45 min at RT in blocking buffer (20% sheep serum (S2350-100, Biowest) in PBS, followed by overnight at 4°C with primary antibodies and fluorescent phalloidin diluted in PBS with 10% sheep serum, then another overnight with secondary antibodies and Hoechst 33342 (H3570, Invitrogen/Thermo Fischer Scientific, 1:2000), rinsed and kept in PBS. Fixation and immunostaining of cells and groups of cells (on coverslips) were similarly, but with shorter times (10 min fixation, 10 min permeabilization, 1-2 hrs incubation with antibodies, all at RT), and coverslips were mounted using antifade mounting media (Slowfade, S36972, Molecular Probes/Thermo Fischer Scientific). For cell-surface labelling, Triton X100 permeabilization after fixation was skipped, but samples were post-fixed after primary antibody incubation, and permeabilized before incubation with the secondary antibodies.

### Confocal microscopy

z-stacks (0.2 to 0.4μm distance between planes) of immunolabelled cells and groups of cells were acquired using an inverted scanning confocal microscope (Leica SP5-SMD), 63x oil objective (1.4NA). Spheroids were imaged using a Nikon inverted microscope coupled to the Andor Dragonfly spinning disk, 40X water objective (1.15NA) as z-stacks of 4 by 4 tile-scans (1μm distance between planes), which where stitched to generate full images of whole spheroids.

### Image quantification

All quantifications were performed using ImageJ. In all cases, cytoplasmic signal was taken as background, and subtracted from the corresponding intensity values. For measurement of relative E-Cadherin and pMLC signal intensities of whole-mount labelled spheroids, the membrane signal was obtained from horizontal confocal slices through thresholding after background subtraction. Values were normalized for each experiment to the average signal in control spheroids. For small groups of cells, peak intensities for pMLC and E-cadherin were measured by drawing a line across two adjacent cells, perpendicular to the cell edges, and running the corresponding plot profile function throughout the z stack to select the planes with respectively highest E-cadherin and pMLC peaks at cell-cell contact, and highest pMLC peaks at each free edge. These measurements were repeated for each cell pair of a group. Relative intensities were calculated from the areas of these peaks after background subtraction. Vinculin FA quantification was based on thresholding and measurement of average intensity and FA total surface per cell (or cell group). Integrated intensity was calculated as vinculin average intensity x surface. Normalization to cell size was obtained by dividing by the maximal horizontal cell area. EpCAM and Trop2 relative signals in cell groups were measured as average intensity on line scans drawn along the plasma membrane at defined locations, either on horizontal sections or on z projections produced using the “reslice” function. The detailed locations in MCF7 cells are detailed in Fig.S4. The same quantification based on manual drawing of lines along cell interfaces was also used for all the data on HCC1500 et MDA-MD-453 cells.

### Quantification of cell morphology

#### MCF7 single cells

Dissociated cells laid on collagen gel were stained with phalloidin and Hoechst and imaged by confocal microscopy. Quantification of cell area was made on maximal horizontal projection, cell height on maximal orthogonal projection after stack reslicing, using ImageJ software.

#### HCC1550 single cells and groups

Z confocal stacks were resliced, and maximal projections from four orientations were used to evaluate angles made between the edge and the ventral interface (acute, roughly right or weakly acute, obtuse). Multiple combinations were compiled into four categories listed in Fig.S5.

#### Traction force microscopy

Stock solution for soft polyacrylamide substrates of 5 kPa rigidity containing far-red fluorescent nanobeads (Bangs laboratory, FC02F, 0.19 μm) were prepared by mixing acrylamide 40% (A4058, Sigma) and bis-acrylamide 2% (M1533, Sigma) in DPBS 1X (PBS without Ca & Mg, CS1PBS01-01, Eurobio Scientific) according to documented relative concentrations (Tse and Engler, 2010; Vignaud et al., 2014). The thin polyacrylamide-based substrate was polymerized between two different glass coverslips (#631-0162, diameter 32 millimeters, thickness No. 1, VWR) prepared as follows. The first coverslip served as base for the gel. It was cleaned using a plasma cleaner machine, then coated with bind-silane (#GE17-1330-01, Sigma) for 3 to 5 min at RT to ensure the attachment of the gel to the coverslip. The second coverslip served to transfer the patterned extracellular matrix. It was also first plasma cleaned, then coated with 0.1 mg/ml PLL-PEG (PLL20K-G35-PEG2K, JenKem Technology) solution for 30 min at RT to obtain a passivated surface. It was then washed with distilled water, dried, then burned for 5 min with UV light through a micropatterning chrome photomask (45 by 45 µm custom-designed H shapes, micropatterned onto chrome photomask by Toppan). This allowed adsorption of the collagen coating at the burned sites resulting in a micropatterned coated coverslip. Collagen type-I was added at 0.5mg/ml in 0.02 N acetic acid and left for 45 min at RT. For gel polymerization, 1 μl of 10% ammonium persulfate (A3678, Sigma), 1 μl of TEMED (T9281, Sigma) and 0.35 μl of the above-mentioned nanobeads were added to 165 μl of the acrylamide-bisacrylamide stock solution. 47 μl of this solution were used to put between the two coverslips for polymerization (30 min, RT). Once polymerized, the collagen-coated top coverslip was gently removed, exposing the collagen H micropatterned gel. Cells were plated on this substrate at a density of 0.5 × 10^5^ per coverslip in a culture dish. The medium of each dish was replaced with fresh medium to wash out cells that didn’t adhere to the substrate (this step avoided ending up with cells that were not on the patterns). The dishes were then kept in the incubator overnight to allow cell division to obtain cell doublets on each H micropattern. Cells and the underneath nanobeads were imaged using an epifluorescence inverted microscope (Nikon Ti2-E2) with 40x air objective (1.15NA) and an Orca Flash 4.0 sCMOS camera (Hamamatsu), with temperature set at 37°C. This first image served as stressed (pulled) state of the beads. Then the cells were removed from the patterns using trypsin and another image of the same position was taken serving as unstressed state of the beads. The displacement field analysis was done using a custom-made algorithm based on the combination of particle image velocimetry and single-particle tracking. Traction forces were calculated using Fourier transform traction cytometry with zero-order regularization (Milloud et al., 2017; Sabass et al., 2008). All calculations and image processing were performed using MATLAB software.

#### Production and analysis of free-floating cell doublets

siRNA transfected cells were dissociated by a 10 min incubation at RT with dissociation buffer. The buffer was then removed, 1 ml of complete culture medium was added to the cells, and single dissociated cells were obtained by pipetting. Dissociated cells of two different conditions (with control cells labelled with Hoechst 33342) were gently mixed, transferred to agarose-coated plastic dishes (2% agarose in PBS) in complete culture medium and put in the incubator for 10 to 15 min to allow partial reassociation. This time was empirically determined as sufficient to reach close to maximal cell-cell contact expansion while minimizing formation of larger groups of cells. Reassociated cells were then gently transferred to a glass-bottom dish coated with a thin layer of 2% agarose, in complete culture medium containing a 1:10000 dilution of the membrane dye CellMask deep red (C1004, Molecular Probes/Thermo Fischer Scientific). Cell doublets were imaged by live confocal microscopy using a Dragonfly spinning disk (Andor) mounted on a Nikon inverted microscope with a 20x (0.75 NA/oil) objective, at 37°C and 5% CO_2_. Hoechst and CellMask images were obtained simultaneously using two CCD cameras (EMCCD iXon888 Life Andor). Quantifications of the relative cortical tensions and adhesiveness were done based geometry of cell membranes at the vertices, as previously described (Kashkooli et al., 2021). Homo- and heterotypic doublets were identified based on the presence of Hoechst-positive nuclei.

#### Statistical analysis

All statistical analyses were performed using the Excel real statistics add-in. Unless otherwise stated, experiments were replicated at least three times and comparisons between conditions were done using either one-way ANOVA followed by Tukey-HSD post hoc test, or the non-parametric Kruskal-Wallis Test, followed by Dunn post-hoc test. p values met the following criteria *p < 0.05, **p < 0.01, and ***p < 0.001 and NS, not significant.

## Supporting information

Annex 1

## Acknowledgements

We acknowledge the help of the MRI imaging platform. We thank Marion Lapierre (IRCM Montpellier) for advice and for providing cell lines. This study was supported by ANR grant ANR-14-ACHN-0004–ICM, ANR-21-CE13-0042-01, and a Labex EpiGenMed Chair of excellence ANR grant to F.F., and CIHR grant 130350 to F.F. and P.L.

## Supplemental material

**Annex.** Description of simulation (cellular Potts model).

**Figure S1**. Related to figure 1

**Figure S2.** Related to figures 1 and 2

**Figure S3**. Related to figure 4 and 5

**Figure S4**. Related to figure 6

**Figure S5.** Related to figure 7

**Figure S6.** Related to figure 7

**Figure S7**. Related to figure 7

**Movie 1.** Related to Figure 1

Time lapse of spreading of siCtrl spheroid on fibrillar collagen gel.

**Movie 2.** Related to Figure 1

Time lapse of spreading of siEpCAM spheroid on fibrillar collagen gel.

**Movie 3.** Related to Figure 1

Time lapse of spreading of siTrop2 spheroid on fibrillar collagen gel.

**Movie 4.** Related to Figure 1

Time lapse of spreading of dKD spheroid on fibrillar collagen gel.

**Movie 5.** Related to Figure 5

Time lapse of simulation of spheroid spreading on the matrix substrate (blue) using 3D Cellular Potts Model. 3D view.

**Movie 6.** Related to Figure 5

Simulation of spheroid spreading on the matrix substrate. Top: xz projection. Bottom: xy projection, plane closest to the substrate.

### Box 1. Description of cell spreading and adhesion based on the balance of tensions at interfaces.

**(A)** The capacity of cells to spread on a substrate, be it matrix or other cells, is strongly dependent on contractility of the actomyosin cortex, which can be viewed as analogous to the role of surface tension in the physical process of the “wetting” of a surface by a liquid (Brodland, 2002, Douezan et al, 2011, Amack and Manning, 2012, Winklbauer 2015). The system can similarly be described as the balance of the tensions exerted at various interfaces, namely tension at the matrix (γ_x_), at cell contact (γ_c_), and along the free edges exposed to the medium (γ_m_). γ_x_ and γ_c_ combine various tensions exerted at the corresponding interface. In the case of cell-cell adhesion, for instance, γ_c_ is the sum of the cortical tensions on each sides of the contact minus the so-called “adhesive tension” (see below). For adhesion to matrix, γ_x_ and γ_m_ are balanced by substrate tension γ_s_. Note that it is customary consider the horizontal components γ_s,h_, which is typically the one measured by techniques such as traction force microscopy. Actin polymerization within the advancing protrusion also generates a force γ_p_, which may contribute to the force balance, depending on the degree of the mechanical coupling with the rest of the cytoskeleton. **(B)** Taking the case of cell-cell adhesion, expansion of an adhesive contact λ requires that contact tension γ_c_ is lowered compared the basal cortical tension γ_m_ that would otherwise act on each side of the interface (2 x γ_m_). This is achieved partly by the adhesion tension resulting from the binding of adhesion molecules to their ligands (cadherin-cadherin and integrins-matrix), but mainly through an active downregulation of local actomyosin contractility along the contact interface. This latter process relies on the ability of adhesion molecules to recruit regulators of cytoskeleton remodelling. To highlight this downregulation, we introduce here a single tension vector γ_α_ that includes the ensemble of contributions that decrease γ_c_ (thus 2γ_m_ = γ_c_ - γ_α_). Panels 1-4 illustrate how cell geometry reflects the force balance: As shown in panel (1), a low γ_c_/γ_m_ ratio (high γ_α_) leads to a large contact interface λ, and a wide angle θ. The latter is a direct geometric readout of “adhesiveness” (David et al, 2014). Contacts adapt to stress through a process of reinforcement, involving recruitment of cortical cytoskeleton and of adhesion molecules, and increased linkage of adhesion molecules to the cytoskeleton (Engl et al, 2014, Charras and Yap, 2018). As shown in panel (2), cells can then maintain the same degree of adhesiveness (same angle θ) despite bearing higher tensions. Importantly, this also requires more repression of contractility along the cell-cell interface (increased γ_α_), another expected effect of enhanced cadherin recruitment. Panel (3) shows the situation where higher cortical tension is not compensated by increased γ_α_, resulting in a higher γ_c_/γ_m_ ratio and lower adhesiveness (smaller angle θ). In the last example (4), γ_m_ remains unchanged compared to (1), but γ_α_ is decreased, resulting in relatively high γ_c_, and thus low adhesiveness (low angle θ and shorter contact interface λ. The same principles are applicable to adhesion and spreading to the extracellular matrix, which depend on the balance between γ_x_ and γ_m_, with spreading of cell aggregates controlled by the combination of cell-matrix and cell-cell adhesion, thus by all three tensions (Ryan et al, 2001, Douezan et al, 2011).

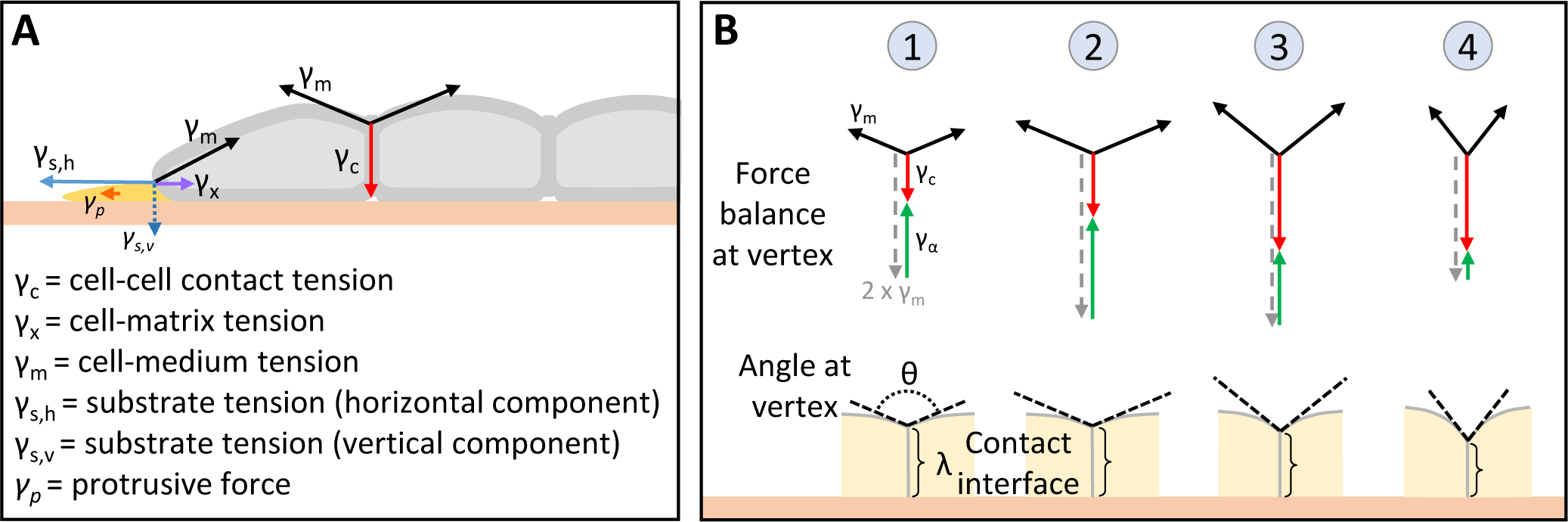

