## Supplementary figures and images for "An EpCAM/Trop2 mechanostat differentially regulates collective behaviour of human carcinoma cells"

### Movie 5

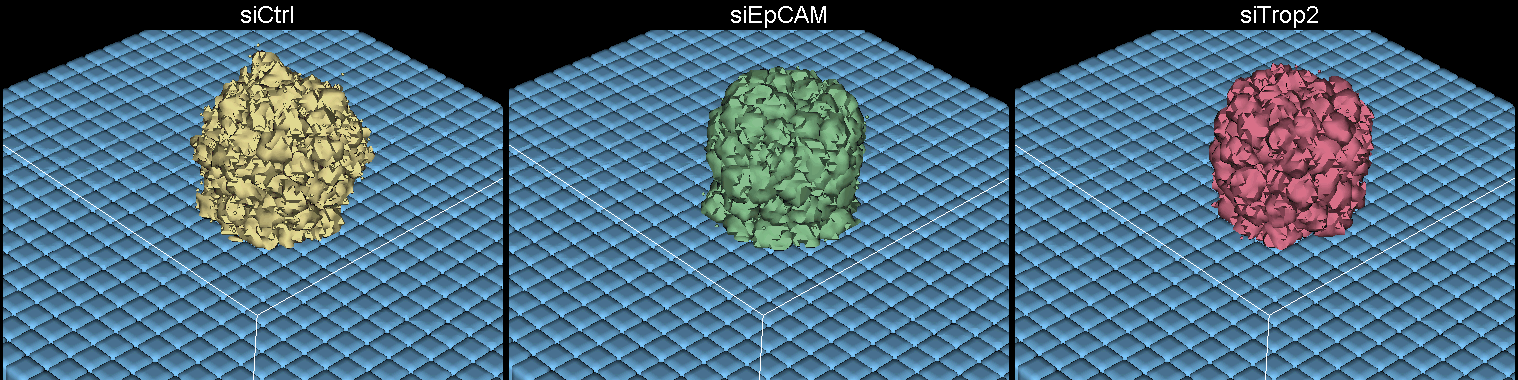

### Movie 6

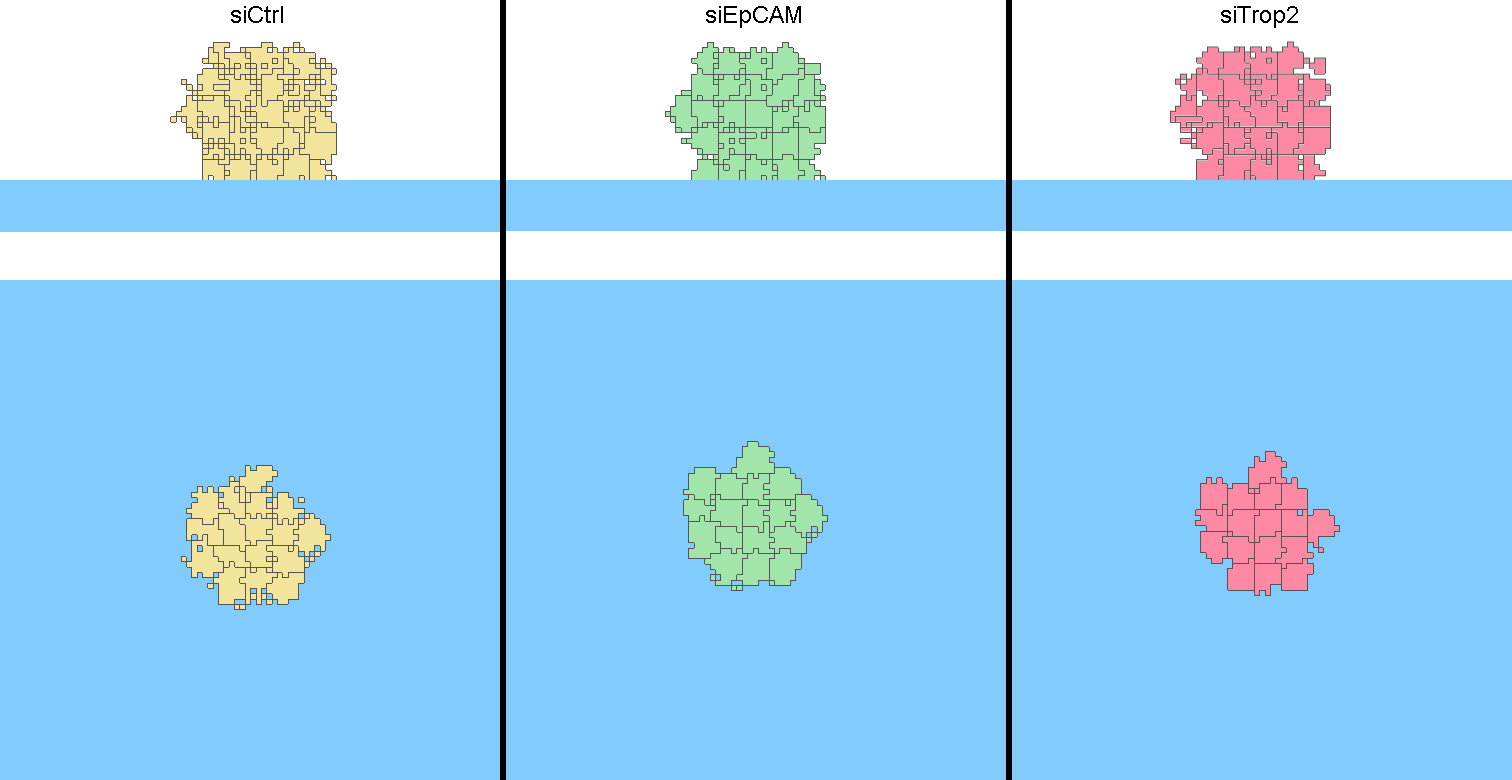
