## Supplemental Figures and legends for "An EpCAM/Trop2 mechanostat differentially regulates collective behaviour of human carcinoma cells"

### Figure S1

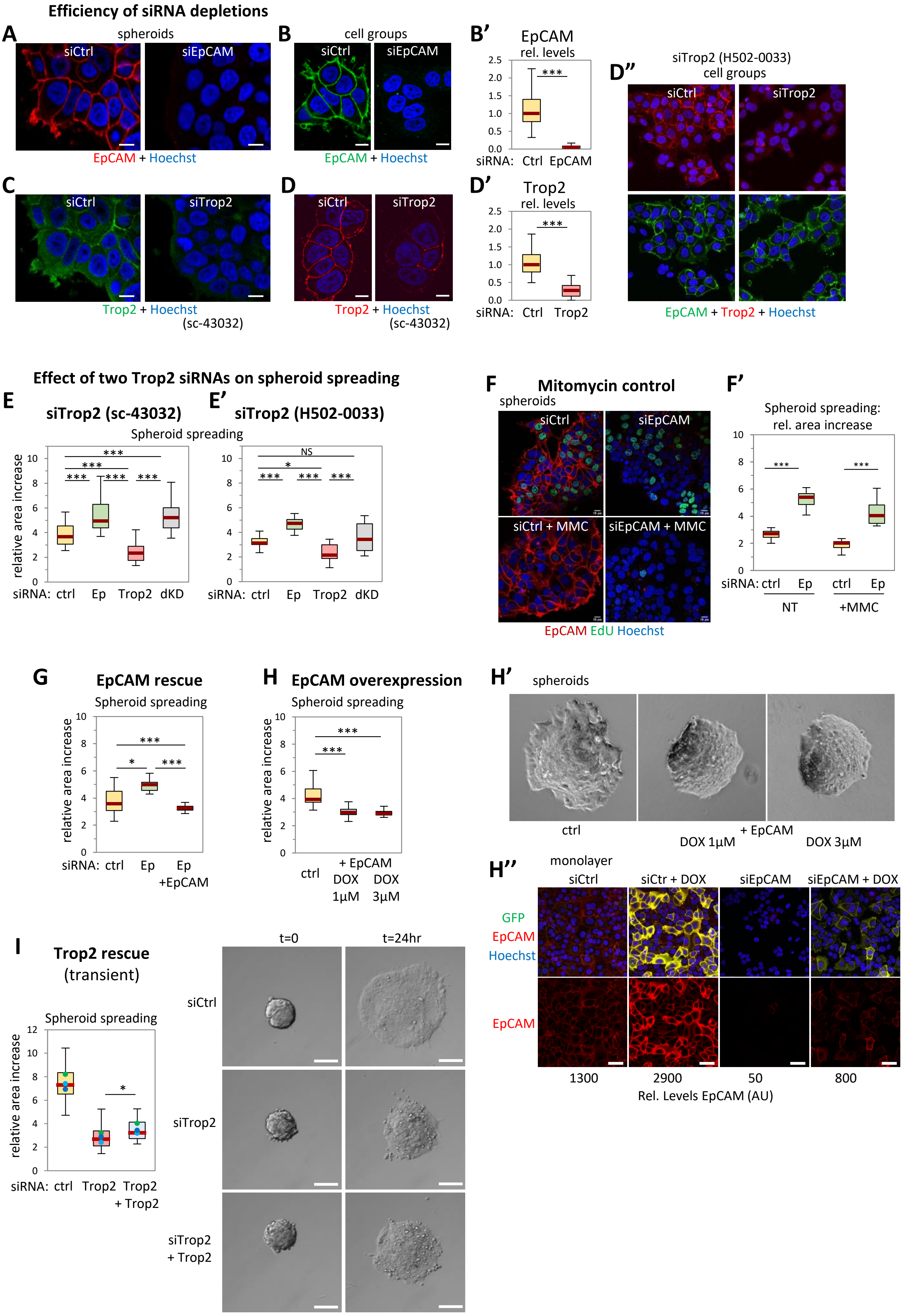

Efficiency of siRNA depletions
A
spheroids
siCtrl
siEpCAM
EpCAM + Hoechst
B
cell groups
siCtrl
siEpCAM
EpCAM + Hoechst
B'
EpCAM
rel. levels
2.5
2.0
1.5
1.0
0.5
0.0
***
siRNA: Ctrl EpCAM
D''
siTrop2 (H502-0033)
cell groups
siCtrl
siTrop2
EpCAM + Trop2 + Hoechst
C
siCtrl
siTrop2
Trop2 + Hoechst
(sc-43032)
D
siCtrl
siTrop2
Trop2 + Hoechst
(sc-43032)
D'
Trop2
rel. levels
2.5
2.0
1.5
1.0
0.5
0.0
***
siRNA: Ctrl Trop2
Effect of two Trop2 siRNAs on spheroid spreading
E
siTrop2 (sc-43032)
E'
siTrop2 (H502-0033)
Spheroid spreading
relative area increase
10
8
6
4
2
0
***
***
***
***
***
siRNA: ctrl Ep Trop2 dKD
NS
*
***
***
***
siRNA: ctrl Ep Trop2 dKD
F
Mitomycin control
spheroids
siCtrl
siEpCAM
siCtrl + MMC
siEpCAM + MMC
EpCAM EdU Hoechst
F'
Spheroid spreading:
rel. area increase
10
8
6
4
2
0
***
***
siRNA: ctrl Ep ctrl Ep
NT
+MMC
G
EpCAM rescue
Spheroid spreading
relative area increase
10
8
6
4
2
0
***
*
***
siRNA: ctrl Ep Ep
+EpCAM
H
EpCAM overexpression
Spheroid spreading
relative area increase
10
8
6
4
2
0
***
***
ctrl
+ EpCAM
DOX 1µM
DOX 3µM
H'
spheroids
ctrl
+ EpCAM
DOX 1µM
DOX 3µM
H''
monolayer
siCtrl
siCtr + DOX
siEpCAM
siEpCAM + DOX
GFP
EpCAM
Hoechst
EpCAM
1300
2900
50
800
Rel. Levels EpCAM (AU)
I
Trop2 rescue
(transient)
Spheroid spreading
relative area increase
12
10
8
6
4
2
0
*
siRNA: ctrl Trop2 Trop2
+ Trop2
t=0
t=24hr
siCtrl
siTrop2
siTrop2
+ Trop2

**Figure S1.** Related to Fig.1.

**A-D. EpCAM and Trop2 depletions. A,C.** Representative confocal microscopy images of the edge of spheroids formed with cell transfected for 96hrs with Ctrl, EpCAM and Trop2 siRNA (sc-43032), immunolabelled for EpCAM and Trop2. Nuclei were stained with Hoechst (blue). The specific signal along cell membranes is undetectable in the respective siRNA condition. **B,D.** Confocal microscopy images of groups of cells transfected for 96hrs with Ctrl, EpCAM and Trop2 siRNA (sc-43032), immunolabelled for EpCAM and Trop2. **C',D'.** Quantification of EpCAM and Trop2 signal intensity at cell-cell contacts. Levels were normalized to the median value for siCtrl cells. Results of 125-157 individual contacts from respectively 3 and 4 independent experiments. Statistical analysis, Student's t-test.

**E. Comparison of the effect of siTrop2 sc-43032 and H502-0033 on spheroid spreading.** Quantification of 30-44 spheroids from four to seven experiments (Refer to S1 Data. F1E, exp1-3,6-9) (**E**) and 16-18 spheroids from two experiments (Refer to S1 Data, F1E, exp4,5) (**E'**). Statistical analysis: one-way ANOVA followed by Tukey-HSD post hoc test. **E".** Immunofluorescence for EpCAM and Trop2 of groups of control and H502-0033-transfected cells.

**F. Increased spheroid migration upon EpCAM KD is independent of cell proliferation.** Spheroids were treated with 2.5µM mitomycin C (MMC) during the entire migration assay. 12 spheroids per condition, three experiments. Statistical analysis: one-way ANOVA followed by Tukey-HSD post hoc test. **F'.** Validation of mitomycin MMC efficiency by imaging EdU incorporation. At the end of migration assay, the spheroids were incubated for 1hr with thymidine analogue EdU, which efficiently incorporates into newly synthesized DNA. EdU was detected in green (see Material and Methods), while EpCAM was detected by immunofluorescence (red) and nuclei were stained with Hoechst (blue). The four panels show representative confocal microscopy images of non-treated and MMC treated spheroids of siCtrl and siEpCAM conditions. Scale bar: 15µm.

**G. Rescue of EpCAM KD spheroid phenotype.** Rescue of spheroid spreading phenotype (24hrs) was performed using a mixed population of MCF7 cells stably transfected with a doxycycline (DOX)-inducible EpCAM-GFP variant that included two conservative point mutations within the siRNA target sequence. Ctrl, transfected with siCtrl, no DOX; siEpCAM, no DOX; siEpCAM + 1µM DOX. Under these conditions, control and siEpCAM spheroids behaved similarly to spread to a similar extend as regular control and EpCAM KD spheroids (Fig.1A). DOX treatment fully rescued spreading to control levels. Results from 20 spheroids per conditions, two independent experiments.

**H. EpCAM overexpression inhibits spheroid spreading.** Spheroids of cells expressing doxycycline-inducible EpCAM-GFP were let spreading for 24hrs on collagen gel. Conditions included non-treated control spheroids, and spheroids treated with 1µM and 3µM DOX. Results of 22 spheroids, from four independent experiments. Statistical analysis: one-way ANOVA followed by Tukey-HSD post hoc test. **H'.** Representative examples for the three conditions, 24hrs spreading. **H''.** Representative EpCAM immunofluorescence images of control, DOX-induced, EpCAM-depleted (siEpCAM) and rescued (siEpCAM + DOX) cells. Cells were double labelled for EpCAM, GFP (to detect exogenous, DOX-induced EpCAM). Total relative EpCAM levels in each image are indicated on the right. Scale bars, 50µm.

**I. Transient expression of Trop2-GFP partially rescues the Trop2 KD spheroid phenotype.** Spheroids of MCF7 cells transfected consecutively siTrop2 and a DOX-inducible Trop2-GFP construct, were laid on collagen gel and left to spread in the absence or in the presence of 0.5-3µM DOX. Positive controls were transfected with siCtrl. Note that spheroids were much smaller than in the other experiments, being formed 100 rather than 400 cells, and expanded more extensively (~7 folds versus ~4 folds). The reason for this protocol modification was that larger spheroids were more severely damaged by cell death caused by the transient transfection of the Trop2-GFP plasmid, independently of DOX induction. Results from 22-27 spheroids from three independent experiments. Statistical comparison, Student's t-test. (D') Examples of control, Trop2 KD (siTrop2 + Trop2-GFP without DOX) and rescue ((siTrop2 + Trop2-GFP with 3µM DOX). Scale bars, 100µm.

### Figure S2

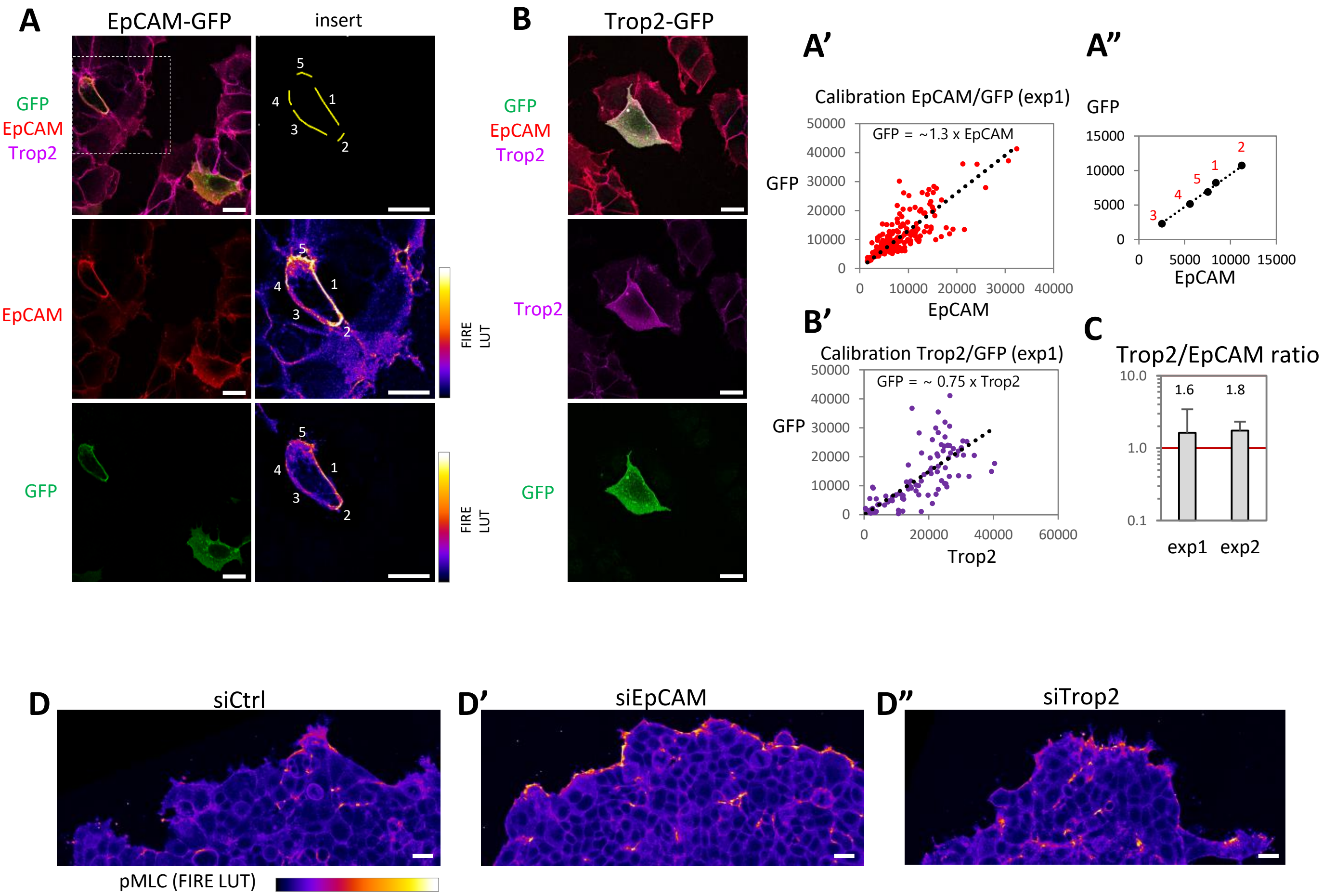


**Figure S2. A-C. Estimate of relative endogenous EpCAM and Trop2 levels in MCF7 cells.** Related to Fig1. The strategy was to use the GFP signal of overexpressed EpCAM-GFP and Trop2-GFP to calibrate the respective signals of the anti-EpCAM and anti-GFP antibodies. Using these calibrations, we could calculate an estimated ratio of endogenous EpCAM and Trop2 in double stained cells. MCF7 cells were transfected with DOX-inducible EpCAM-GFP and Trop2-GFP constructs, treated overnight with 1μM DOX, fixed and immunolabelled, without permeabilization, for EpCAM and Trop2, then postfixed and permeabilized, and immunolabelled for GFP (positioned at the cytoplasmic C-terminus). **(A,B)** Examples of confocal images, showing individual transfected cells among wild type MCF7 cells. **(Insert)** Example of measurement on one EpCAM-GFP positive cell: 6 membrane segments of various intensities, numbered 1-6, were used to quantify GFP and EpCAM signal intensities plotted in A". **(A',B')** Compilation of these values for experiment 1 (6 images for each condition), which were used to determine the approximative slopes. **(C)** These slopes allowed to directly compare endogenous EpCAM and Trop2 levels based on the signal intensities in non-transfected cells. Results from 2 experiments, 10-12 fields. Average values are indicated on a log scale. Bars: SD. This is a rough estimate, as levels strongly varied between cells, even though the GFP to EpCAM/Trop2 signal ratio within individual cells often resulted quite linear (A"). Scale bars: 20μm. **D. Detail of pMLC signal at the periphery of control, EpCAM KD and Trop2 KD spheroids.** Selected planes just above the collagen surface (the broadest area of each spheroid). Related to figure 2. Scale bars, 10μm.

### Figure S3

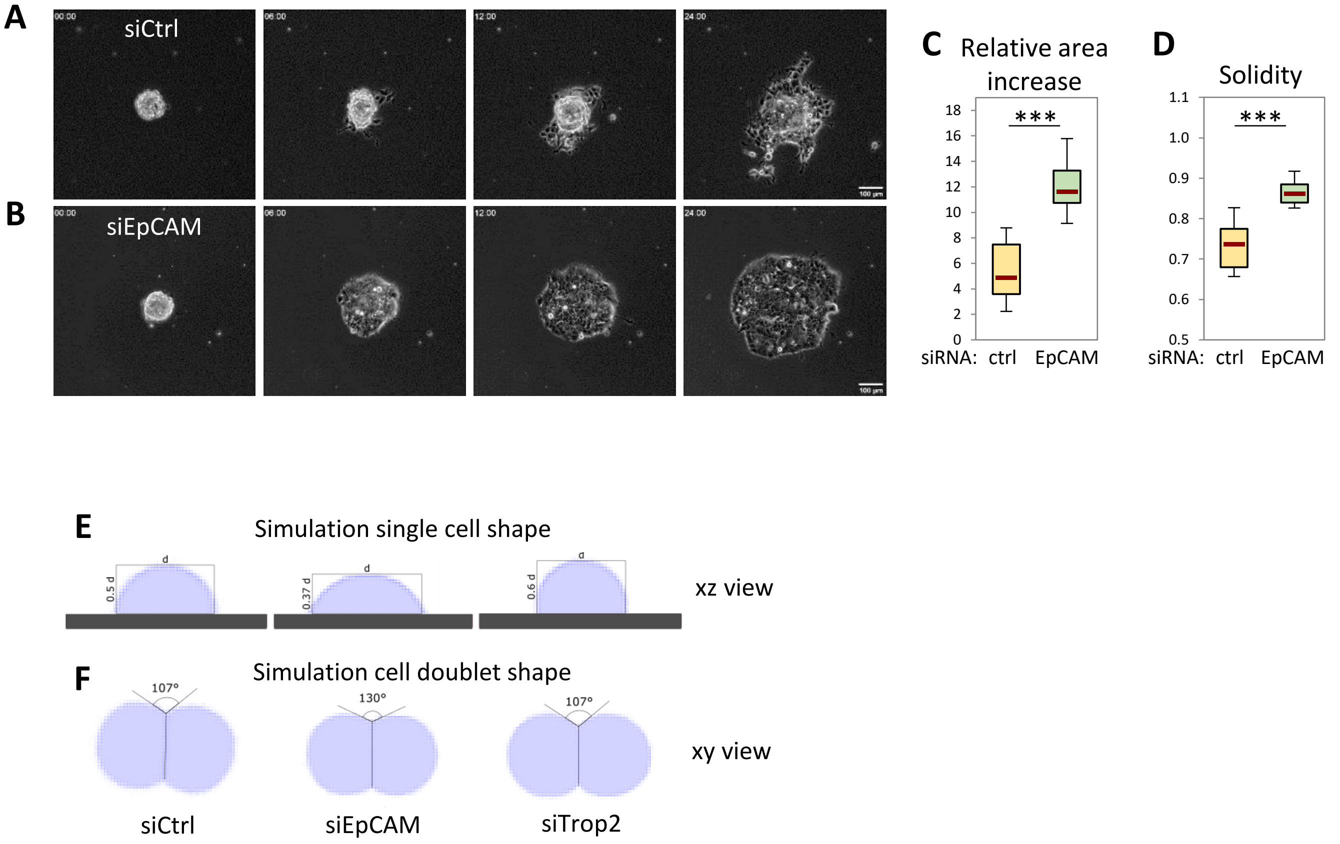


**A-D. Spheroid spreading on collagen-coated 5kPa polyacrylamide gel.** Related to Fig. 4.
**A,B.** Images of siCtrl and siEpCAM spheroids at four time points as in Figure 1. EpCAM KD spheroids spread much more than controls. Control spheroids are very irregular, with cells at the edge bulging out and even detaching. EpCAM KD spheroids remain much more coherent. Scale bars: 150µm.
**C,D.** Quantification of area increase and solidity, as in main Fig.1. 22 spheroids per conditions from four independent experiments. Statistical analysis: Student's t-test.
**E,F. Cellular Potts Model simulation of basic morphology of single cells and doublets based on experimental data.** Related to Fig. 5. **E.** Profile view of single cell geometry obtained using CompuCell3D simulation and the same energy parameters used for simulation of spheroids spreading (Fig. 5). The numbers correspond to the height/diameter ratio (assuming a circular cell surface). Such dimensions approximate those estimated from experimental morphological measurements (Fig. 1I). **F.** Top view of simulated non-adhesive cell doublets, which approximate the experimental data, in particular the average contact angles (Fig. 4H,L).

### Figure S4

#### Analysis of EpCAM and Trop2 cell surface distribution in MCF7 cells

**A** **A’**

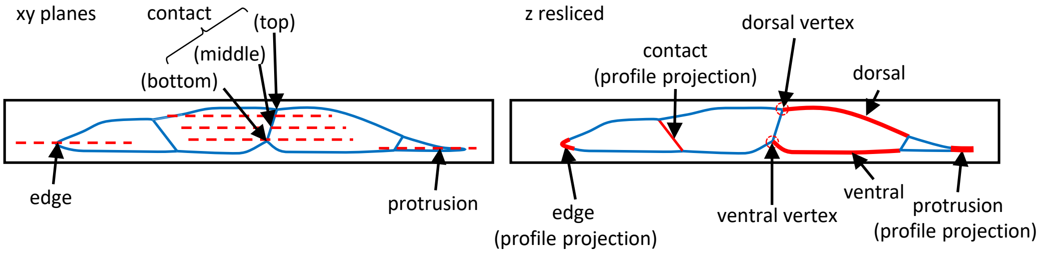


**B** Distribution of EpCAM and Trop2 in cell surface regions of MCF7 cells (Relative levels normalized to ave contacts)

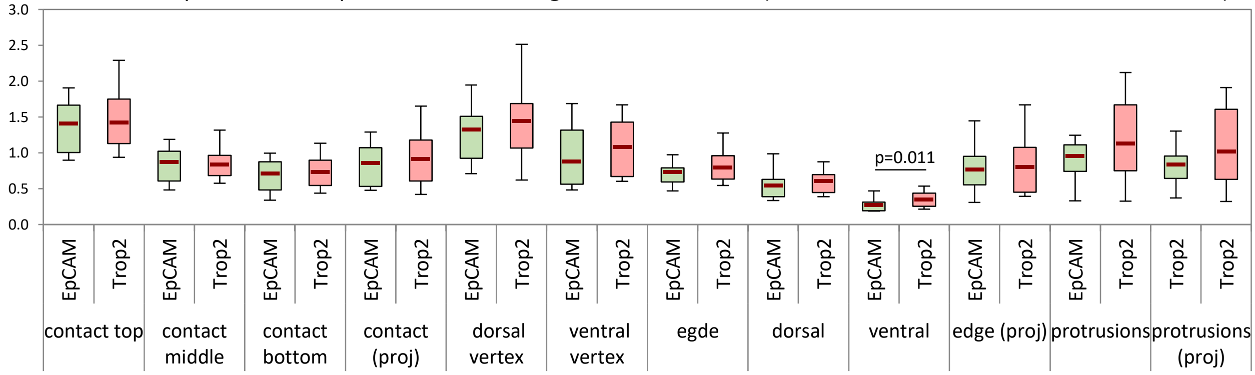


**C** Impact of EpCAM or Trop2 KD on distribution of the other component (Relative levels normalized to ave contacts)

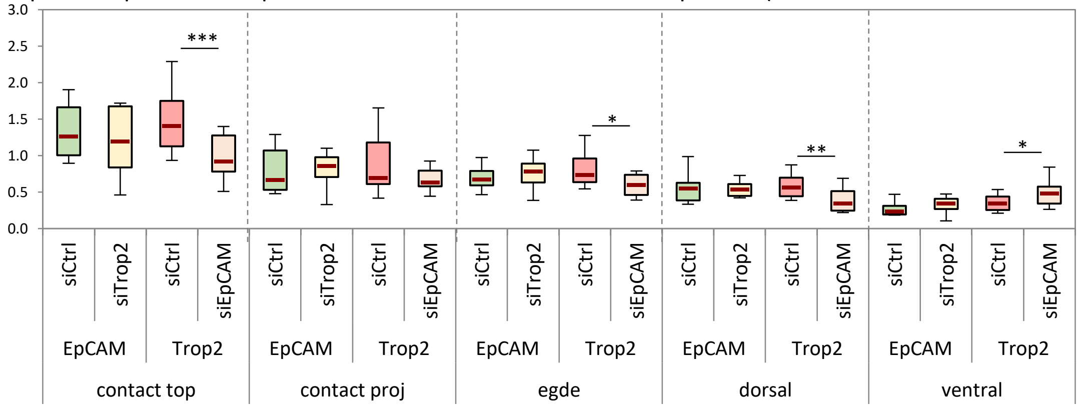


**D** EpCAM/Trop2 distribution in large group of MCF7 cells (non-permeabilized)

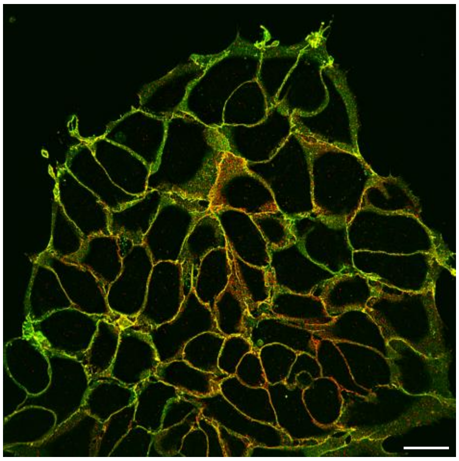

EpCAM Trop2

**E** EpCAM/Trop2 distribution (post-fixation permeabilization)

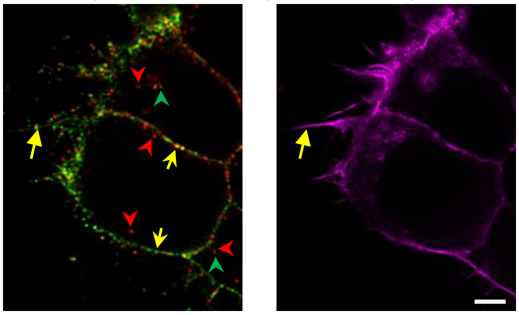

EpCAM Trop2 F-actin

**Figure S4. Detailed analysis of EpCAM and Trop2 distribution.** Related to Figure 6.

**A.** Schematic representation of a group of cells, viewed in profile, showing the typical position of horizontal (xy) planes used to measure line fluorescence intensities along edge and contact cell membrane. **A'.** Same scheme, showing regions measured in profile projections, obtained by reslicing the image stacks. Intensities were also measured from lines, except for vertices, for which small circles were used. A total of 12 subcellular regions were measured. Values of each experiment were normalized to the average value from of the three regions of cell-cell contacts in xy sections, taken near the top, middle and bottom of each contact (bracket in A).

**B.** Quantification of EpCAM and Trop2 levels in control MCF7 cells. Note that several of the categories corresponded to the overlapping regions, measured either in the original horizontal planes or in the corresponding profile projections. The two types of measurements show high consistency, e.g. cell edges and cell protrusions (also compare average contact in xy planes versus whole contacts in profile projections). EpCAM and Trop2 showed quasi identical distributions. The only statistically significant difference was found along the ventral membrane. 13-15 groups of cells (4-8 cells per group) from three independent experiments. Statistical analysis: Non-parametric ANOVA (Kruskal-Wallis Test) followed by post-hoc Pairwise Mann-Whitney tests.

**C.** Effect of EpCAM depletions on Trop2 levels and, reciprocally, of Trop2 depletion on EpCAM levels, for selected regions of the cell membrane. Neither EpCAM levels nor its distribution are significantly changed by Trop2 depletions. Upon EpCAM KD, Trop2 distribution is modified at four localization: Trop2 levels drop at lateral edges and dorsal side, including the dorsal region of cell-cell contacts (contact top), but not average levels along the whole cell-cell contact (profile). Trop2 levels are prominently increased at the ventral side. Student's t-test between control and corresponding KD condition. Only statistically significant pairs are indicated. Pairwise Student's t-test.

**D.** Example of large cell group showing a typical enrichment of Trop2 along internal contacts compared to EpCAM. Maximal projection of 7 planes, total thickness 1.5µm. Scale bar: 20µm.

**E.** Example of immunofluorescence of EpCAM and Trop2 in cells permeabilized with 0.2% Triton X100 after fixation. Both signals along the plasma membrane (yellow arrows) are more punctate compared to those obtained without permeabilization (Fig 8 and panel F). Yellow filled arrows: Punctate signal at protrusions. Trop2 signal at the plasma membrane was much lower than using surface labelling without permeabilization. Few intracellular spots positive for EpCAM or Trop2 are detected (green and red arrowheads). Scale bar: 5µm.

### Figure S5

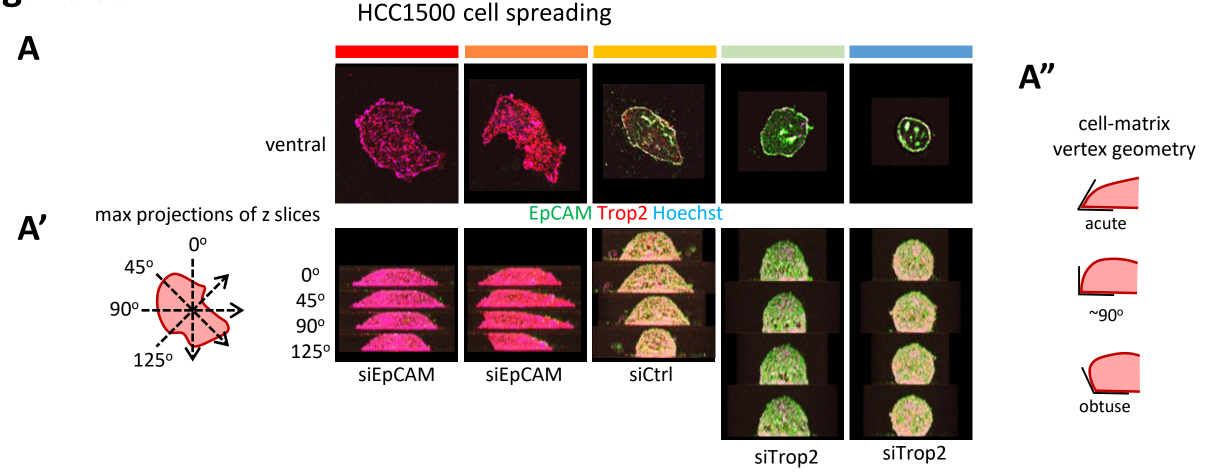


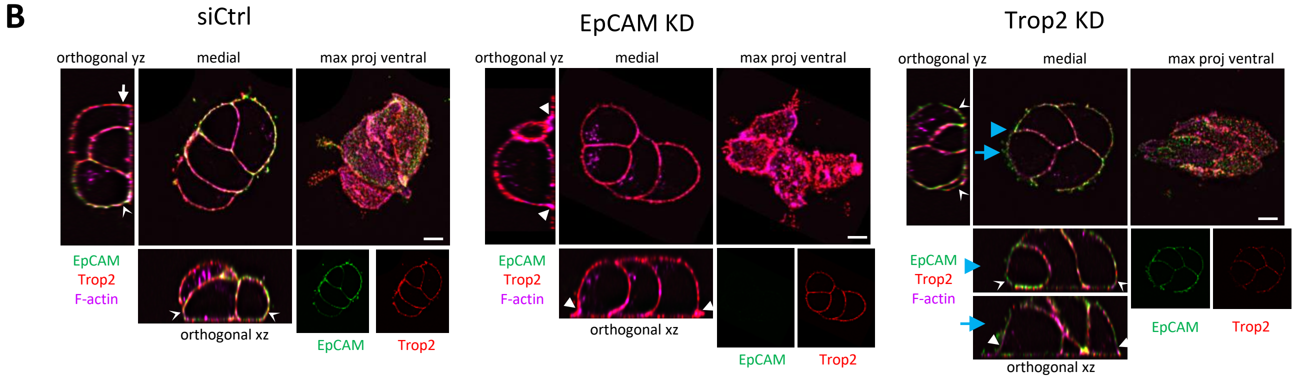


**C**

| Category | | Score | Combined edge geometries from four profiles (8 edges) |
|---|---|---|---|
| Spread | Full | 6 | All acute |
| | High | 4 | Most acute |
| | Moderate | 2 | All ~90º or weakly acute<br>Half ~90º, half obtuse<br>Part obtuse, part acute<br>Half acute, half ~90º |
| | Weak | 1 | Most obtuse, rest ~90º or weakly acute |
| Round | | 0 | All obtuse |

**D**

| P values | SC | Doublets | Groups |
|---|---|---|---|
| siK-siEpCAM | <0.001 | NS | 0.006 |
| siK-siTrop2 | NS | 0.022 | NS |
| siEpCAM-siTrop2 | <0.001 | 0.002 | 0.004 |

HCC1500 spheroids

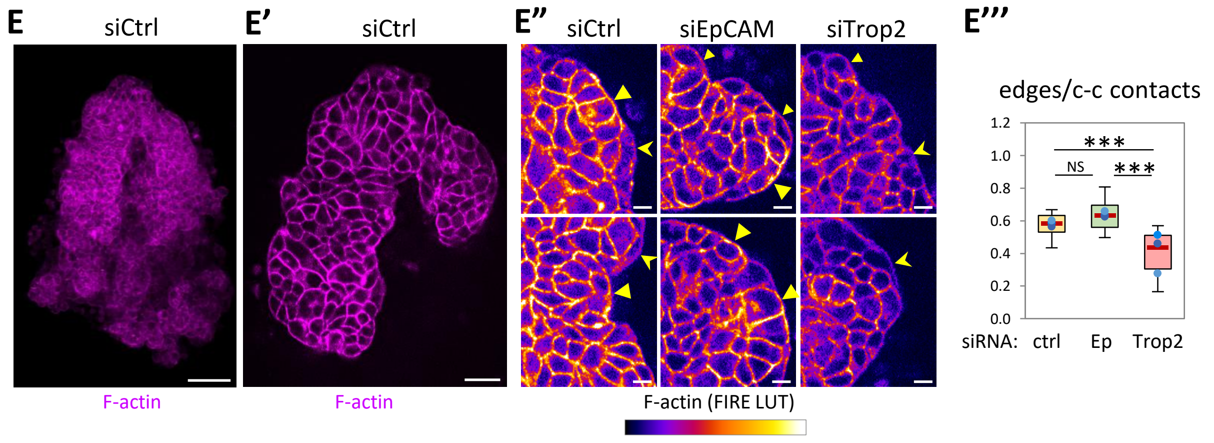

**Figure S5. EpCAM KD and Trop2 KD phenotypes for HCC1500 cells.** Related to Figure 7.

**(A-D) Analysis of morphology phenotypes for single cells (A) and small groups (B).** HCC1500 cells transfected with siCtrl, siEpCAM or siTrop2, laid on fibrillar collagen, were surface immunolabelled for EpCAM, Trop2 and stained for F-actin (Phalloidin). **(A)** Representative examples of single cells, selected to cover the range of degree of spreading. **(A')** Orthogonal maximal projections in four orientations. **(A")** These projections were used to observe the geometry at the contact with the matrix substrate, specifically the angle formed by the free cell edge and the substrate interface. **(B)** Examples of siCtrl, siEpCAM and siTrop2 groups of cells. Each multiple panel includes merged images of a medial horizontal plane, a maximal projection of bottom (ventral) planes, and two orthogonal views xz and xy. Arrowheads point to the free-edge substrate vertex. Filled arrowhead: acute angle; arrow: right angle; convex arrowhead: obtuse angle. For the Trop2 KD example, two xz slices are shown, one with obtuse acute angles (blue arrowhead), one with acute angles (blue arrow). Scale bars: 5µm. Separate EpCAM and Trop2 channels of the median plane are shown as small inserts for visualization of their respective levels under control and depletion conditions. **(C)** Morphological classification. Single cells and cell groups were classified in five categories of degree of spreading, from round to fully spread, based on the geometry at the substrate vertex. Because of the highly irregular shapes, multiple combinations were pooled in the intermediate categories. **(D)** Statistical analysis corresponding to the results shown in the main figure 7C. P values from ANOVA analysis followed by Tukey-HSD post-hoc test, obtained by allocating to each categories a score (0,1,2,4,6). Results were essentially the same using different scales (e.g. 0,1,2,3,4). **(E) EpCAM KD and Trop2 KD differentially impact on the cortex at free edges and cell-cell contact in HCC1500 spheroids.** Spheroids formed from control, siEpCAM and siTrop2 cells and laid on collagen gel for 3 days were stained with phalloidin. (E) Overview of a control spheroid, presented as maximal projection. HCC1500 spheroids are typically highly convoluted. (E') Single slice from the same spheroid. (E") Detailed views of spheroid edges, two examples per condition. Arrowheads point to free edges, to be compared with cell-cell contacts: Large filled, small filled and concave arrowheads indicate respectively high, moderate and low F-actin. (E"') Quantification of the edge/contact intensity ratio. 20-24 spheroids from 3 experiments. Dots indicate averages for each experiment. One-way non-parametric ANOVA (Kruskal-Wallis Test) followed by Dunn post-hoc test. Scale bars: B, 100µm; B', 50µm; C, 10µm.

### Figure S6

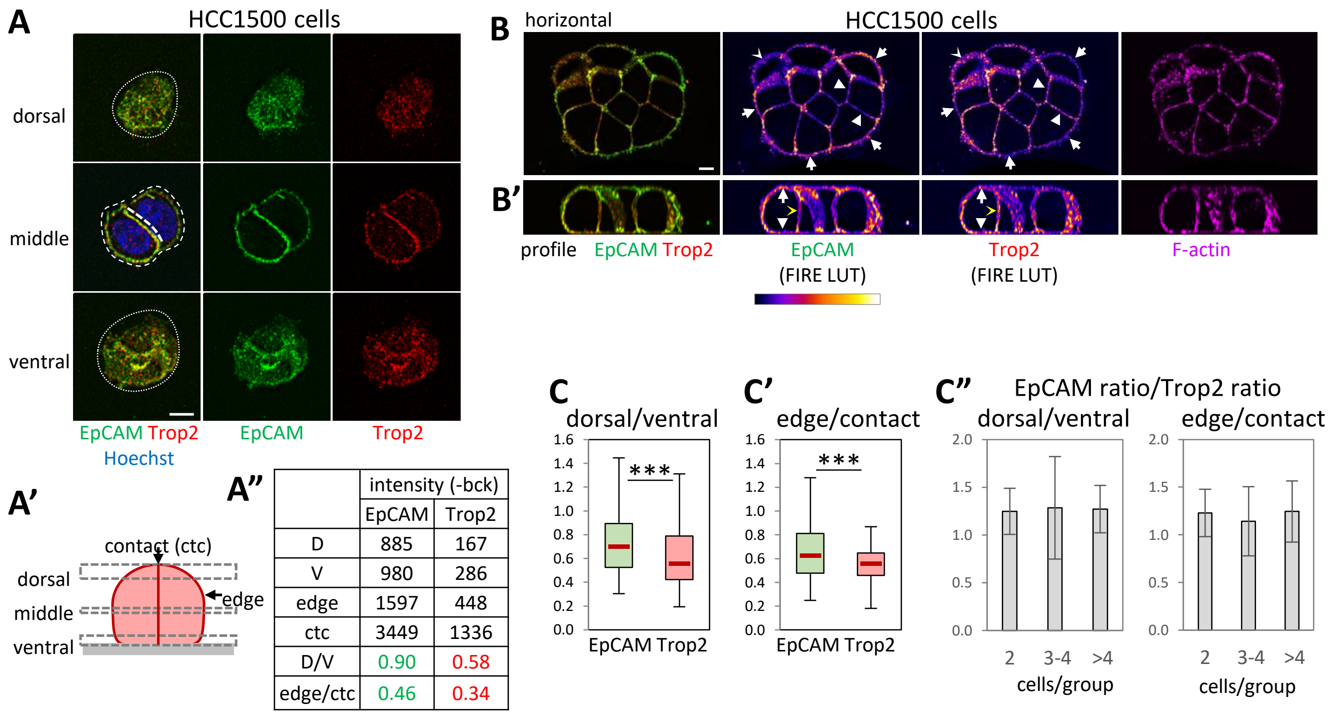


| | intensity (-bck) | |
|---|---|---|
| | EpCAM | Trop2 |
| D | 885 | 167 |
| V | 980 | 286 |
| edge | 1597 | 448 |
| ctc | 3449 | 1336 |
| D/V | 0.90 | 0.58 |
| edge/ctc | 0.46 | 0.34 |

**Figure S6. (A-C) Differential EpCAM and Trop2 distribution in HCC1500 cells.** Related to Fig.7. HCC1500 cells were surface immunolabelled for EpCAM and Trop2, and stained for Hoechst (A) or phalloidin (B). **(A)** Cell doublet. Three horizontal slices corresponding to dorsal (0.2µm max projection, 2 planes), middle (0.8µm max projection, 5 planes) and ventral (0.2µm max projection, 2 planes) regions (A'). Dotted lines indicate the dorsal and ventral areas (horizontal plane), and dashed lines the free edge and cell contact interfaces (line scans on z profiles) used for quantification. Scale bar, 5µm. **(A")** Quantification for this doublet, providing average intensities, background subtracted, for dorsal and ventral surfaces, and for the lateral edge and cell-cell contact (ctc), and the resulting dorsal/ventral (D/V) and ctc/edge ratios. **(B)** Group of cells labelled for EpCAM, Trop2 and F-actin (phalloidin). Arrows point to edges with higher EpCAM relative to Trop2, arrowheads to contacts with low EpCAM and higher Trop2. Small concave arrowhead: Case of edge with high Trop2. **(B')** Orthogonal view, used for quantification. White arrow points to dorsal edge, with high EpCAM, white arrowhead to the ventral interface, and yellow concave arrowhead to cell contact, both with higher Trop2. **(C,C')** Quantification of dorsal/ventral and edge/contact ratios for EpCAM and Trop2 intensities, for groups ranging from doublets up to 20 cell-large groups. **(C")** Calculated EpCAM ratios divided by Trop2 ratios for groups of different sizes, showing that the relative enrichments are constant over the whole range. 67 groups from 3 experiments. Statistical analysis using Student's t-test. Scale bar: 20µm.

### Figure S7

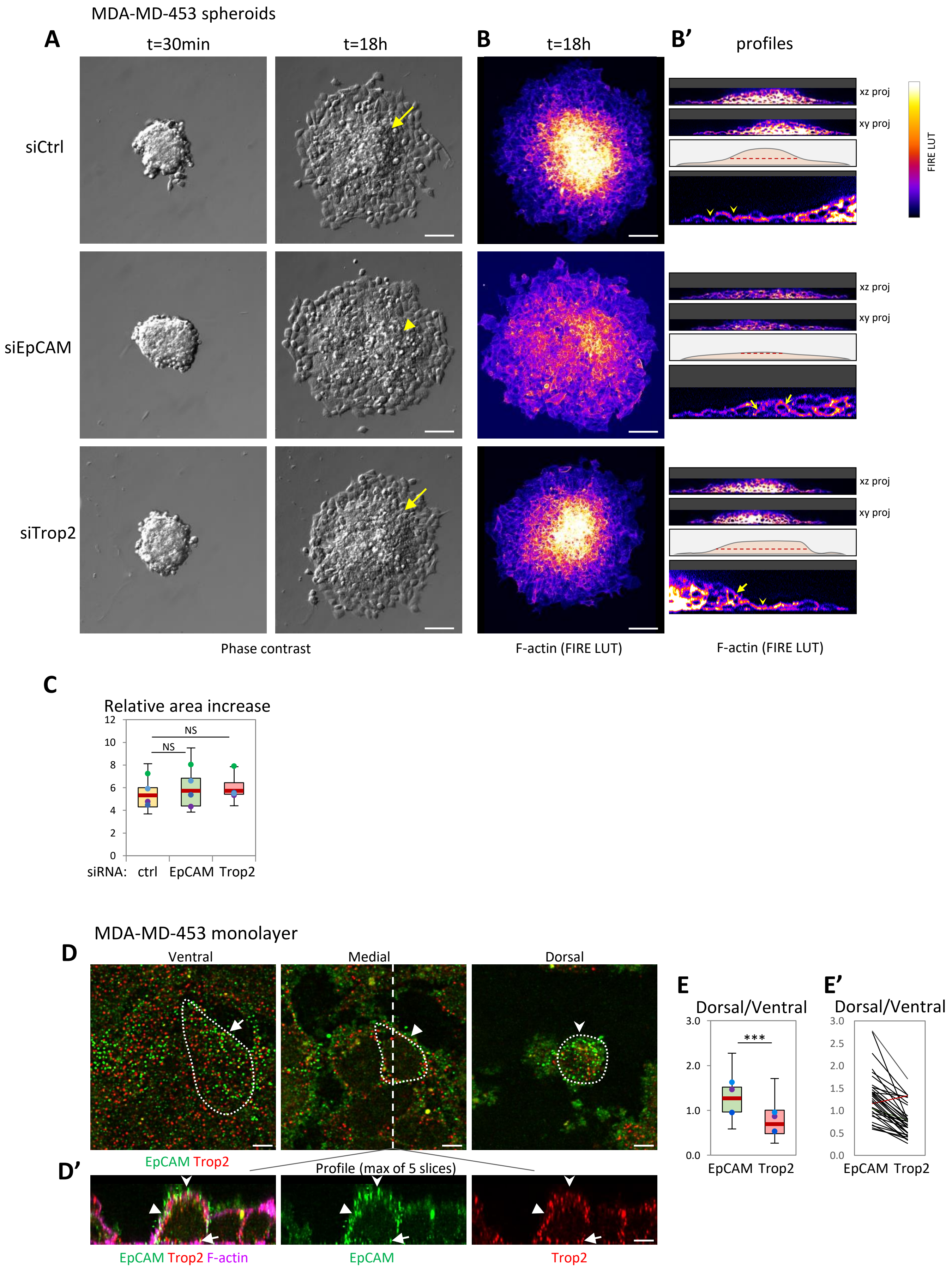

MDA-MD-453 spheroids
A
t=30min
t=18h
B
t=18h
B'
profiles
siCtrl
siEpCAM
siTrop2
xz proj
xy proj
FIRE LUT
Phase contrast
F-actin (FIRE LUT)
F-actin (FIRE LUT)
C
Relative area increase
NS
NS
0
2
4
6
8
10
12
siRNA: ctrl EpCAM Trop2
MDA-MD-453 monolayer
D
Ventral
Medial
Dorsal
EpCAM Trop2
D'
Profile (max of 5 slices)
EpCAM Trop2 F-actin
EpCAM
Trop2
E
Dorsal/Ventral
***
0.0
1.0
2.0
3.0
EpCAM Trop2
E'
Dorsal/Ventral
0.0
0.5
1.0
1.5
2.0
2.5
3.0
EpCAM Trop2

**Figure S7. (A) Spreading of MDA-MD-453 spheroids.** Related to Fig.7. Bright field images of control, EpCAM KD and Trop2 KD spheroids, shortly after adhering to the collagen gel (t=30min), and after 18hrs. At 18hrs, spheroids have extensively spread. In control spheroids, a remnant core of the original cell mass can still be seen as a dome (arrow). In EpCAM KD spheroids, the dome is much smaller (arrowhead) and often absent. In Trop2 KD spheroids, the core is typically larger than in controls. **(B)** Maximal projection of spread spheroids labelled for F-actin with Phalloidin, shown as FIRE LUT. **(B')** Profiles obtained from xz and yz orthogonal projections. Top panels show the two profiles of the spheroid, bottom panel an enlarged view. The middle diagram schematizes the characteristic morphology for each of the three conditions. The red dashed line indicates the emerging dome. **(C)** Quantification of the area increase, was not significantly different between conditions. 30-33 spheroids from 4 experiments. One-way non-parametric ANOVA (Kruskal-Wallis Test) followed by Dunn post-hoc test. **(D-E) EpCAM and Trop2 localization in wild type MDA-MD-453 cells**. Cells were densely plated on a collagen gel. EpCAM and Trop2 signals were variable and generally low. **(D)** Slices of the ventral, medial and dorsal planes are shown, with the outlines of one cell indicated by a dotted line. **(D')** Profile obtained from orthogonal slices, maximum projection of 5 slices. Arrow: ventral interface; filled arrowhead: lateral surface; concave arrowhead: dorsal side. **(E,E')** Dorsal/ventral ratio of EpCAM and Trop2 intensities shows that EpCAM is enriched dorsally and Trop2 ventrally. Quantification of 33 cells from 3 experiments. Dots: Averages of individual experiments. E' shows the pair of ratios for each cell. Red line indicates the only cell where the ratios were inverted. Statistical analysis using Student's t-test. Scale bars: A,B, 50μm; D, 5μm.
