## Supplementary material for "An EpCAM/Trop2 mechanostat differentially regulates collective behaviour of human carcinoma cells": Annex 1

### Annex – CompuCell3D Simulations

All Simulations were performed with CompuCell3D, a software package that simulates cell behavior in a 3D environment using a Cellular Potts Model. For a detailed description of the model, see the corresponding publication (Maciej H. Swat, 2012).

In short, the model consists of a regular, 3D lattice where each pixel belongs to a specific, user-defined generic cell. These cells can correspond to actual cells, the substrate, the medium or other components and the user can assign attributes to these cells and their interactions with other cells. Next, the user specifies an initial configuration, which then evolves over time according to the user-defined interaction rules.

The evolution of the initial configuration proceeds by minimizing the overall energy in the system. More concretely, in each simulation step, some pixels change their state randomly, where the number of pixels that change per step is defined with the *temperature* parameter, and the overall energy of the new configuration is calculated. If the energy of the new configuration is smaller than the energy of the previous configuration, the new configuration is accepted. If it is larger, it can also be accepted with a certain probability, dependent on a so-called Hamiltonian function, which inputs parameters, including target cell area/volume and interaction between cell types. In the present simulations, there were three different types of cells: the substrate, the medium and the cells. The interaction between each pair of cell types is defined by their corresponding binding energies. This yielded 6 different energy parameters:

- Medium-medium energy
- Medium-cell energy
- Medium-substrate energy
- Cell-cell energy
- Cell-substrate energy
- Substrate-substrate energy

Since both the medium and the substrate are immobile, i.e. they never switch state unless replaced by a cell, their interactions with themselves and with each other don't impact the result of the simulation. In other words, the medium-medium and the substrate-substrate energy parameter are irrelevant. From the four remaining parameters, only three are independent. Since the system only tries to minimize the overall energy by comparing different states, the total energy of the system is arbitrary. Therefore, we can set one of these parameters to 0 without losing any degrees of freedom. We chose to set the medium-substrate energy to 0, which leaves us with three remaining free parameters: The cell-cell, the cell-substrate and the cell-medium energy.

Choosing these three parameters has the advantage that they all correspond to concrete physical properties of the cells. The cell-medium energy corresponds to the cell cortical tension $\gamma_m$, the cell-cell energy corresponds to contact tension $\gamma_c$, which is inversely related to cell-cell adhesiveness, and the cell-substrate energy corresponds cell substrate tension $\gamma_x$, also inversely related to cell-substrate adhesiveness.

This direct correspondence allowed us to identify the parameters for the different conditions one by one. First, we set $\gamma_m$ for ctrl cells $\gamma_{m(C)}$ to 2. Based on global contractile energies obtained from TFM data, we set $\gamma_m$ for siEpCAM cells $\gamma_{m(E)}$ twice higher, i.e. to 4, and that for siTrop2 cells $\gamma_{m(T)}$ slightly lower, i.e. 3.5. Starting with these values, cell-substrate energies were estimated based on the geometry of single cells laid on collagen (S8 Fig): We calculated a theoretical diameter from the area measurements in S8 Fig, by assuming a circular spreading area, according to:

$$A = \pi\left(\frac{d}{2}\right)^2 => d = 2\sqrt{\frac{A}{\pi}}$$

We then calculated the median height to diameter ratio for the three different conditions, and then simulated single, spreading cells, where we varied the cell-substrate energy until the height-width ratio corresponded to the single cell spreading data from. To deal with the stochastic and pixelated nature of the simulation, we calculated an average image over 200 simulation time points. For control cells, cell-substrate energy was 0, consistent with a 0.5 ratio, thus forming a half circle. (S8 Fig). The cell-substrate energy for siEpCAM (ratio 0.37) was -2. For siTrop2, we needed to slightly deviate from the median measurement of 0.7 to better capture the spheroid spreading phenotype observed for this condition, and set the energy to +1 (ratio 0.6).

We set the last free parameter, cell-cell energy based on simulation of non-adherent cell doublets, akin to the experimental data shown in Figure 6H-L. Just like for the single spreading cells, we varied the cell-cell energy parameter, until the simulated doublets resembled the doublets from the experiments. With this approach we fixed the cell-cell energy to 3 for siCtrl, 5 for siEpCAM and 6 for siTrop2. Note that this is only an approximation, since it is very difficult to measure a contact angle on these pixelated, averaged images. These energy parameters are summarized in Figure 7G. We used these parameters to simulate spreading spheroids made of approximately 100 cells.
